# Transglutaminase mediated asprosin oligomerization allows its tissue storage as fibers

**DOI:** 10.1101/2022.01.04.474899

**Authors:** Yousef A.T. Morcos, Galyna Pryymachuk, Steffen Lütke, Antje Gerken, Alan R. F. Godwin, Thomas A. Jowitt, Nadin Piekarek, Thorben Hoffmann, Anja Niehoff, Margarete Odenthal, Uta Drebber, Olaf Grisk, Yury Ladilov, Wilhelm Bloch, Bert Callewaert, Mats Paulsson, Eva Hucklenbruch-Rother, Clair Baldock, Gerhard Sengle

## Abstract

Asprosin, the C-terminal furin cleavage product of profibrillin-1, was reported to act as a hormone that circulates at nanomolar levels and is recruited to the liver where it induces G protein-coupled activation of the cAMP-PKA pathway and stimulates rapid glucose release into the circulation. Although derived upon C-terminal cleavage of fibrillin-1, a multidomain extracellular matrix glycoprotein with a ubiquitous distribution in connective tissues, little is known about the mechanisms controlling the bioavailability of asprosin in tissues. In the current view, asprosin is mainly produced by white adipose tissue from where it is released into the blood in monomeric form. Here, by employing newly generated specific asprosin antibodies we monitored the distribution pattern of asprosin in human and murine connective tissues such as placenta, and muscle. Thereby we detected the presence of asprosin positive extracellular fibers. Further, by screening established cell lines for asprosin synthesis we found that most cells derived from musculoskeletal tissues render asprosin into an oligomerized form. Our analyses show that asprosin already multimerizes intracellularly, but that stable multimerization via covalent bonds is facilitated by transglutaminase activity. Further, asprosin fiber formation requires an intact fibrillin-1 fiber network for proper linear deposition. Our data suggest a new extracellular storage mechanism of asprosin in an oligomerized form which may regulate its cellular bioavailability in tissues.

## Introduction

Fibrillin-1 is a large, 350 kDa extracellular matrix (ECM) glycoprotein that when mutated leads to the multi-system disorder Marfan syndrome (MFS; OMIM**#**154700) affecting mostly the musculoskeletal (e.g. long bone overgrowth, muscle wasting, hyperflexible joints) and cardiovascular system (e.g. aortic root aneurysm formation). The multiple clinical features of MFS reflect the ubiquitous tissue distribution pattern of fibrillin-1 (1). Since most mutations in *FBN1* result in a slender habitus with little subcutaneous fat (2), it is intuitive to speculate that fibrillin-1 plays a role in metabolism and body fat formation. In addition, specific mutations within exon 64 at the 3’ end of *FBN1* were described to lead to marfanoid-progeroid-lipodystrophy syndrome (MFLS; OMIM**#**616914) (3), a fibrillinopathy characterized by progeroid facial features and severe lipodystrophy (3-6). A mechanistic explanation for these striking clinical features was provided by the discovery that the C-terminal cleavage product of profibrillin-1 serves as a fasting-induced glucogenic protein hormone that modulates hepatic glucose release (7). Profibrillin-1 is translated as a 2,871-amino-acid long proprotein, which is cleaved at the C-terminus by the protease furin (8). In addition to mature fibrillin-1, this cleavage generates a 140-amino-acid long C-terminal cleavage product. MFLS mutations were found to be clustered around the furin cleavage site, thereby causing heterozygous ablation of the C-terminal cleavage product which was only detectable at strongly reduced levels in plasma of patients (7). This C-terminal furin cleavage product of fibrillin-1 was termed “asprosin” after the Greek word for white because of its strong impact on white fat tissue (WAT) (7).

Asprosin was reported to circulate at nanomolar levels and is recruited to the liver where it induces G protein-coupled activation of the cAMP-PKA pathway and stimulates rapid glucose release into the circulation (7). Following a circadian rhythm and triggered by fasting, asprosin induces hepatic glucose release via the G-protein-coupled OLFR734 receptor (7,9) and reduces insulin secretion from pancreatic β-cells (10). Asprosin has also been shown to cross the blood brain barrier and activate hunger-stimulating AgRP (Agouti-related peptide) neurons in the hypothalamus which induces appetite in mice (11). In addition, asprosin was shown to impair insulin sensitivity in skeletal muscle cells *in vitro* (12). There are also several reports of increased serum asprosin levels in obese patients and patients with type 2 diabetes mellitus (T2DM) and that asprosin serum concentrations are positively correlated with insulin resistance (13). In mice, antibody-mediated neutralization of asprosin leads to reduced food intake and body weight as well as improved insulin sensitivity (11). Therefore, asprosin currently ranges among the most promising candidates for the pharmacological therapy of obesity, T2DM and other metabolic disorders (13).

Although derived from ubiquitously expressed fibrillin-1, little is known about the tissue distribution of asprosin. In the current view, asprosin is mainly produced by WAT from where it is released into the blood in monomeric form. However, our previous findings showed that asprosin is synthesized and secreted by connective tissue resident cells such as fibroblasts and chondrocytes (14) – a finding which supports the notion of local asprosin storage within tissue-specific microenvironments. Here, by employing newly generated specific asprosin antibodies we monitored the distribution pattern of asprosin in various human and murine connective tissues, including placenta, and muscle. In addition, we monitored the synthesis of asprosin in various established cell lines. Our data reveal new insight into how asprosin may be stored within the extracellular microenvironment of connective tissues and how this affects its cellular bioavailability.

## Results

### Extracellular storage of asprosin as fibers in tissues

Since fibrillin-1 shows a broad tissue distribution, we wanted to monitor asprosin expression and its potential intra- or extracellular storage by tissues. Recently, we were able to generate specific anti-asprosin antibodies against human and mouse asprosin that allow the sensitive detection of asprosin in clinical samples and do not cross-react with fibrillin-1 (14).

Here, we show that also our newly generated antibody against mouse asprosin does not crossreact with human asprosin (supplementary Fig. S1). Therefore, we employed both newly generated antibodies to monitor the tissue distribution of asprosin in human and murine tissues (Fig. 1). Asprosin signals were detected as slight diffuse or vesicle like staining in hepatocytes which can be recognized by their large round nuclei (Fig. 1F, L). In skeletal and cardiac myocytes, anti-asprosin antibodies showed a crossstriated appearance with higher intensity on intercalation discs of cardiomyocytes (Fig. 1C, J). In addition, asprosin positive signals were also detected between myocytes in the form of thin fibers (Fig. 1B, I) In cryosections of cruciate ligament (Fig. 1E) and heart valve biopsies (Fig. 1D, K), asprosin positive fibers were even more pronounced. In the kidney, asprosin fibers were also observed in the glomeruli (Fig. 1G, M). Also, in human placenta asprosin positive fibers were observed (Fig.1A), which were less prominent in mouse placenta (Fig.1H), likely due to species specific differences in the placenta ECM architecture (15).

**Figure 1:**
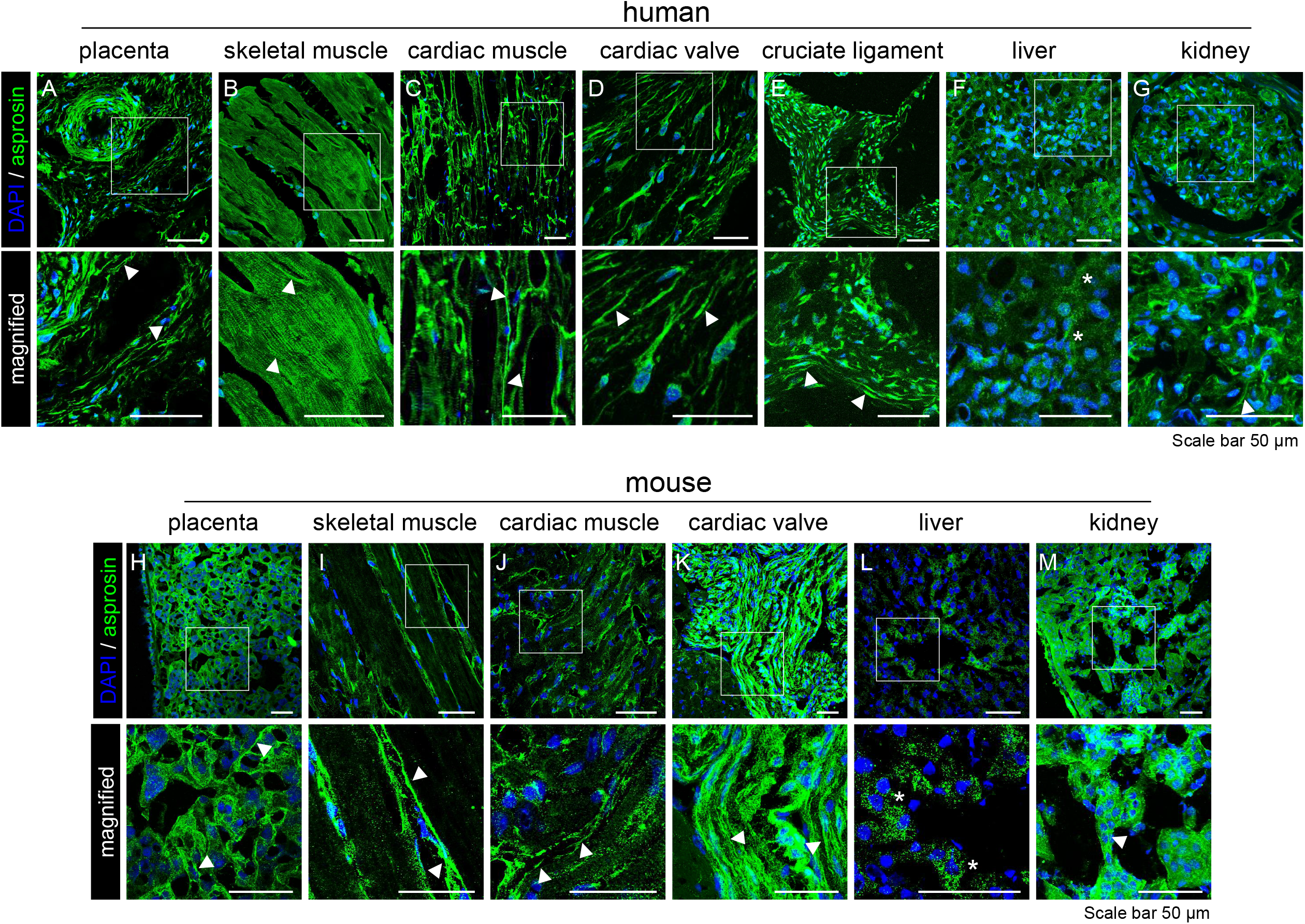
Detection of asprosin positive fibers in human and mouse tissues. Distribution of asprosin in human (A-F) and mouse (G-L) tissues. Cryosections were fixed with acetone and incubated with anti-human or anti-mouse asprosin antibody (green) and DAPI (blue, nuclei). Extra- and intracellular signals of endogenous asprosin were mainly detected in two staining patterns: fiber-like structures (white arrow heads) and diffused or versicle-like structures (asterisks). 2 to 3-fold magnified areas are marked by white boxes Images were obtained from a Leica SP8 confocal microscope and were processed using Leica Application Suite X (LAS X) software version 3.7.5.2. Fiji/ImageJ (version 1.53t) software was used to obtain average intensity Z-projection.

### Asprosin is detected as multimers in cultures of cell lines

To investigate asprosin synthesis and secretion by cells derived from connective tissues, we screened different cell lines for intra- and extracellular presence of asprosin by immunoblot analysis. We detected asprosin positive bands of higher molecular weight suggesting the presence of asprosin multimers (Fig. 2A). Thereby we observed a higher abundance of asprosin multimers in cell culture supernatants of cell lines derived from musculoskeletal tissues such as U2OS (bone osteosarcoma epithelial cells) or C2C12 (mouse myoblasts) when compared to WI-26 (embryonic lung fibroblasts) or RPE (retinal pigment epithelial cells) (Fig. 2A). To investigate the extracellular deposition of asprosin in cell cultures, we employed immunofluorescence analysis after 7-9 days of culture time. Interestingly, similar to our findings in tissues we also detected asprosin positive fibers within the ECM deposited by certain cell lines such as 3T3-L1 fibroblasts (model for fibro/adipogenic progenitors present in skeletal muscle) as well as WI-26, primary human skin fibroblasts (HDF) and human chondrocytes (HCH) (Fig. 2B).

**Figure 2:**
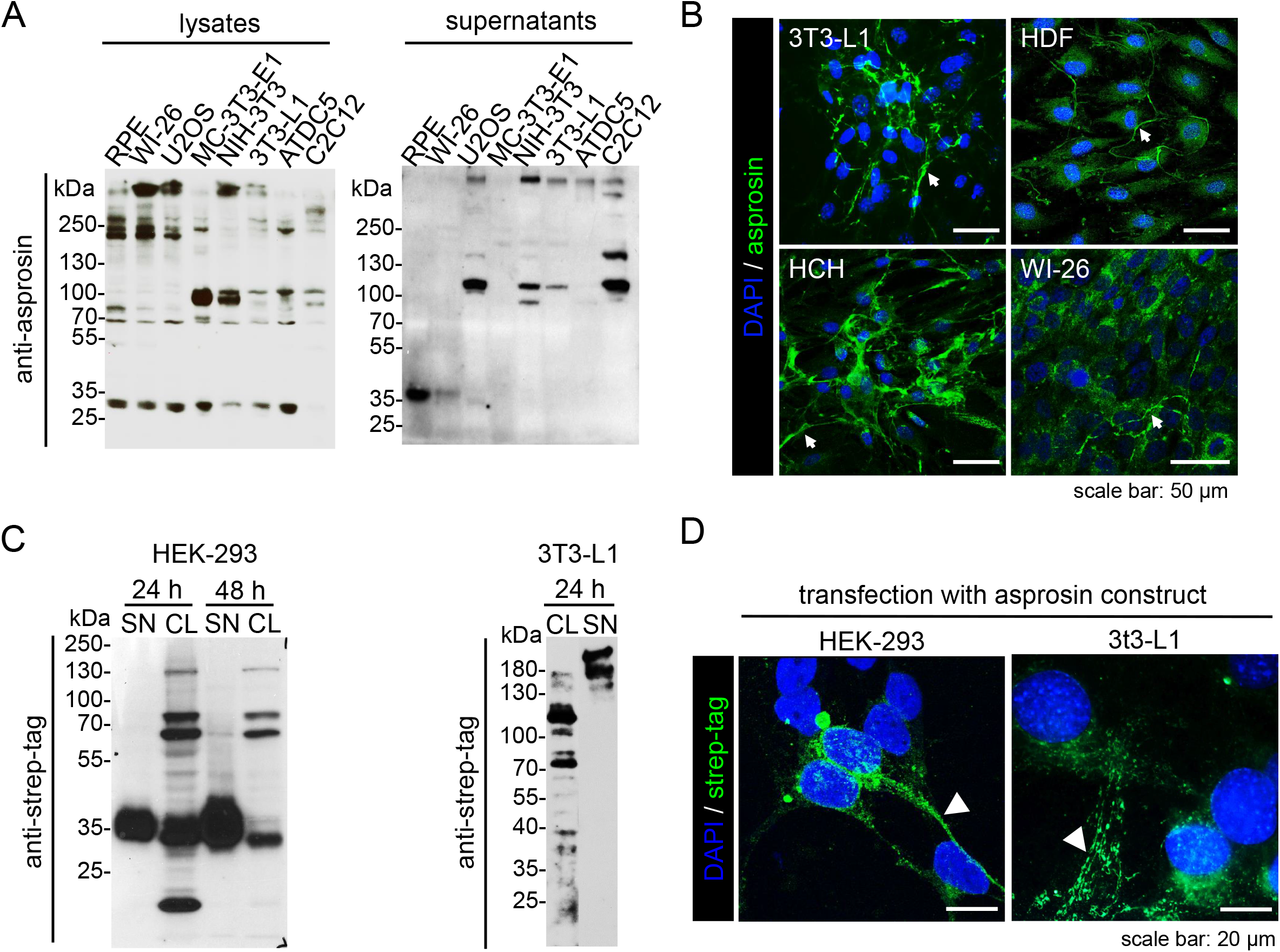
Asprosin multimers and fibers detected in cell culture and upon overexpression of monomeric asprosin. **A**. Western blot analysis of asprosin in cell lysates and cell culture from different cell lines. Asprosin was detected at the corresponding molecular weight of ∼37 kDa and as bands of higher molecular weight in both cell lysates and supernatants. **B**. Immunofluorescence analysis of asprosin in 3T3-L1; primary human dermal fibroblasts (HDF), and primary human chondrocytes (HCH) after 7-9 days in culture and WI-26 after 3 days in culture. 3T3-L1 cells were incubated with anti-asprosin (mab-Asp Ab) (green) and DAPI (blue, nuclei). HDF, WI-26 and HCH were incubated with anti-asprosin (pc-asp Ab) (green) and DAPI (blue, nuclei). The staining reveals depositions of asprosin positive fibers as indicated by white arrow heads. **C**. Immunoblotting of cell lysates and cell culture supernatants from HEK-293 and 3T3-L1 transfected with pCEP-Pu vector overexpressing asprosin-2×Strep-tag II using anti-streptag (StrepMAB Ab). Immunoblots detected monomeric asprosin (∼37 kDa) bands in the supernatant of transfected HEK-293 cells, but high molecular weight bands in the corresponding cell lysate fraction. However, western blot analysis of supernatants and lysates of transfected 3T3-L1 cells only showed asprosin positive multimer bands. **D**. Immunofluorescence analysis of asprosin-transfected HEK-293 and 3T3-L1 cells with anti-streptag (StrepMAB Ab) showing asprosin intra- and extracellular positive signals. The extracellular signals reveal deposition of asprosin positive fibers as indicated by white arrowheads. Images were obtained from Axiophot Microscope (Carl Zeiss, Germany) and Leica SP8 confocal microscope and were processed using Leica Application Suite X (LAS X) software version 3.7.5.2. Fiji/ImageJ (version 1.53t) software was used to obtain average intensity Z-projection.

### Overexpression of monomeric asprosin in musculoskeletal cells results in multimerization

To further investigate the formation of asprosin multimers in musculoskeletal cells, we transfected 3T3-L1 cells with an overexpression construct encoding for monomeric asprosin. Western blot analysis of the collected conditioned media and cell lysates after SDS-PAGE showed that most asprosin was detected in bands representing higher molecular weight multimers that persisted even under reducing conditions (Fig. 2C). However, when HEK-293 cells were transfected with the same construct, only asprosin monomers were detected in the cell culture supernatant (Fig. 2C). Interestingly, signals representing multimerized forms of asprosin were also detected in HEK-293 cell lysates and were resistant to reducing agents. Immunofluorescence analysis of cultured 3T3-L1 cells showed the presence of fibers consisting of endogenously expressed asprosin (Fig. 2B, top-left) as well as overexpressed monomeric asprosin after transfection (Fig. 2D, right).

Moreover, immunofluorescence analysis of HEK-293 cells stably transfected with human asprosin revealed a fibril like structure of the overexpressed asprosin (Fig. 2D; supplementary Fig. S2B, left-column). Also, similar asprosin positive structures were detected in stably transfected cells overexpressing murine asprosin (supplementary Fig. S2B, middle-column). However, in cells overexpressing placensin, the C-terminal propeptide of fibrillin-2 (16), we detected only intracellular signals with no evidence of fiber formation (supplementary Fig. S2B, right-column) by employing a newly generated placensin antibody (supplementary Fig. S3A-C).

### Intrinsic multimerization ability of recombinant asprosin

When recombinant asprosin was subjected to size exclusion chromatography (SEC), the obtained chromatogram showed two main asprosin containing peaks eluting at fraction numbers corresponding to higher molecular weight species (Fig. 3A) than the previously determined 37 kDa of glycosylated recombinant asprosin (14). In line with this finding, analysis of asprosin fractions directly after affinity purification by native-PAGE revealed the accumulation of asprosin in the stacking gel with no detected migration into the gel (Fig. 3D, right). However, SDS-PAGE analysis of collected fractions from both peaks indicated a prominent monomeric asprosin band at 37 kDa under these denaturing conditions (Fig. 3B). Interestingly, in fractions 14-17 of the second peak an additional band migrating above 50 kDa corresponding to the size of potential asprosin dimers was also detected (Fig. 3B). To determine the size distribution and multimeric status of asprosin, SEC was performed followed by multiangle light scattering (MALS). The MALS profile indicated a polydispersity of asprosin, with a polydispersity index (PDI) of 1.140 ± 5.561% and a molecular mass ranging from ∼180 kDa to ∼1000 kDa (an average of 570 kDa) and average hydrodynamic radius (R_h_) of 11.38 nm (Fig. 3C).

**Figure 3:**
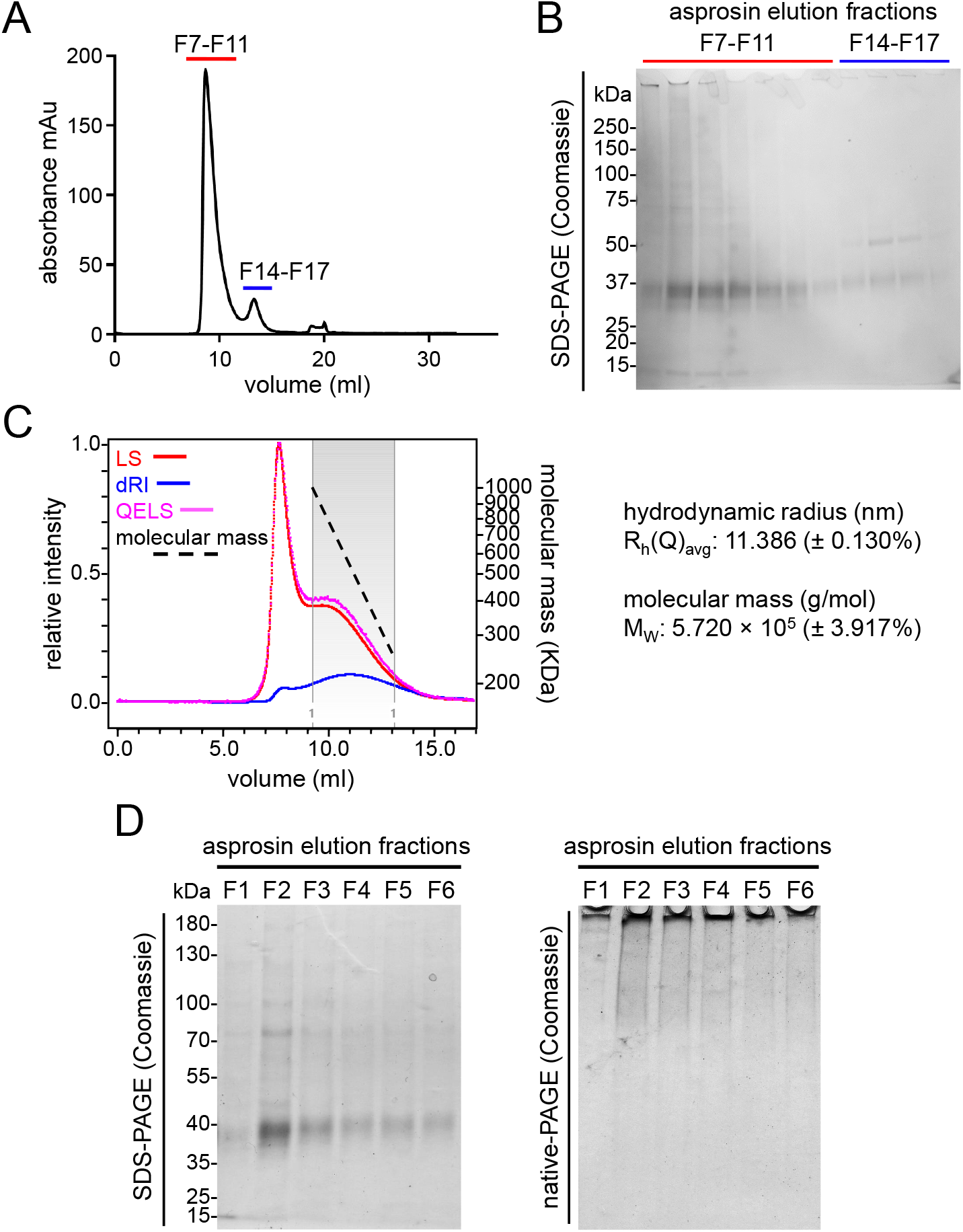
Asprosin forms multimers under native conditions. **A**. Size exclusion chromatogram of asprosin after affinity purification. Most asprosin protein elutes in one peak at 9 ml (F7-F11, marked in red), while a minor amount elutes in F14-F17 (marked in blue). **B**. SDS-PAGE analysis of asprosin containing peak elution fractions shown in A under non-reducing conditions. **C**. MALS analysis after size exclusion chromatography of affinity purified asprosin. Blue line indicates refractive index (RI), red line indicates scattering at 90 degrees, pink line indicates dynamic light scattering (DLS). Dashed black line indicates molecular mass against elution volume, showing high polydispersity ranging from 180 to 1000 kDa (average mass of 570 kDa). **D**. Coomassie stained gels of recombinantly purified asprosin (with C-terminal 2×Strep-tag II). Eluted fractions (F2-F6) after affinity chromatography were subjected to (left) a reducing 10% SDS-PAGE gel and (right) a 10% native-PAGE gel. The Coomassie stained native-PAGE gel shows an accumulation of asprosin multimers in stacking gel and at stacking/running gel interface.

In order to identify conditions in which asprosin multimers form or dissolve, we subjected recombinant asprosin to significant changes in temperature (20°C to 90°C) (supplementary Fig. S4B), pH (3-4, 8-9) or buffers (Urea, DTT). Our findings indicated that asprosin multimers are resistant to the applied changes (supplementary Fig. S5B-G). To further determine which conditions favor or abolish asprosin multimerization, we employed the high-throughput UNcle protein stability analyzer (supplementary Fig. S4B-F). Thereby, we observed a sufficient dissociation of asprosin multimers upon SDS addition, which resulted in significant reduction of the average hydrodynamic radius from ∼117 nm to ∼17 nm upon addition of 2 % SDS (Fig. 4A). Asprosin dissociation upon SDS addition was monitored by electrophoresis showing separation of asprosin multimers and migration into the gel to different positions by native-PAGE analysis (Fig. 4B, middle and right), while SDS-PAGE results remained unchanged (Fig. 4B, left). In addition, since it was reported that polyamines have a preventive effect on protein aggregation (17), we subjected asprosin to spermine and spermidine treatment. Native-PAGE analysis of asprosin after treatment with spermine or spermidine resulted in a dissociation pattern similar to SDS treatment (supplementary Fig. S5 H, I).

**Figure 4:**
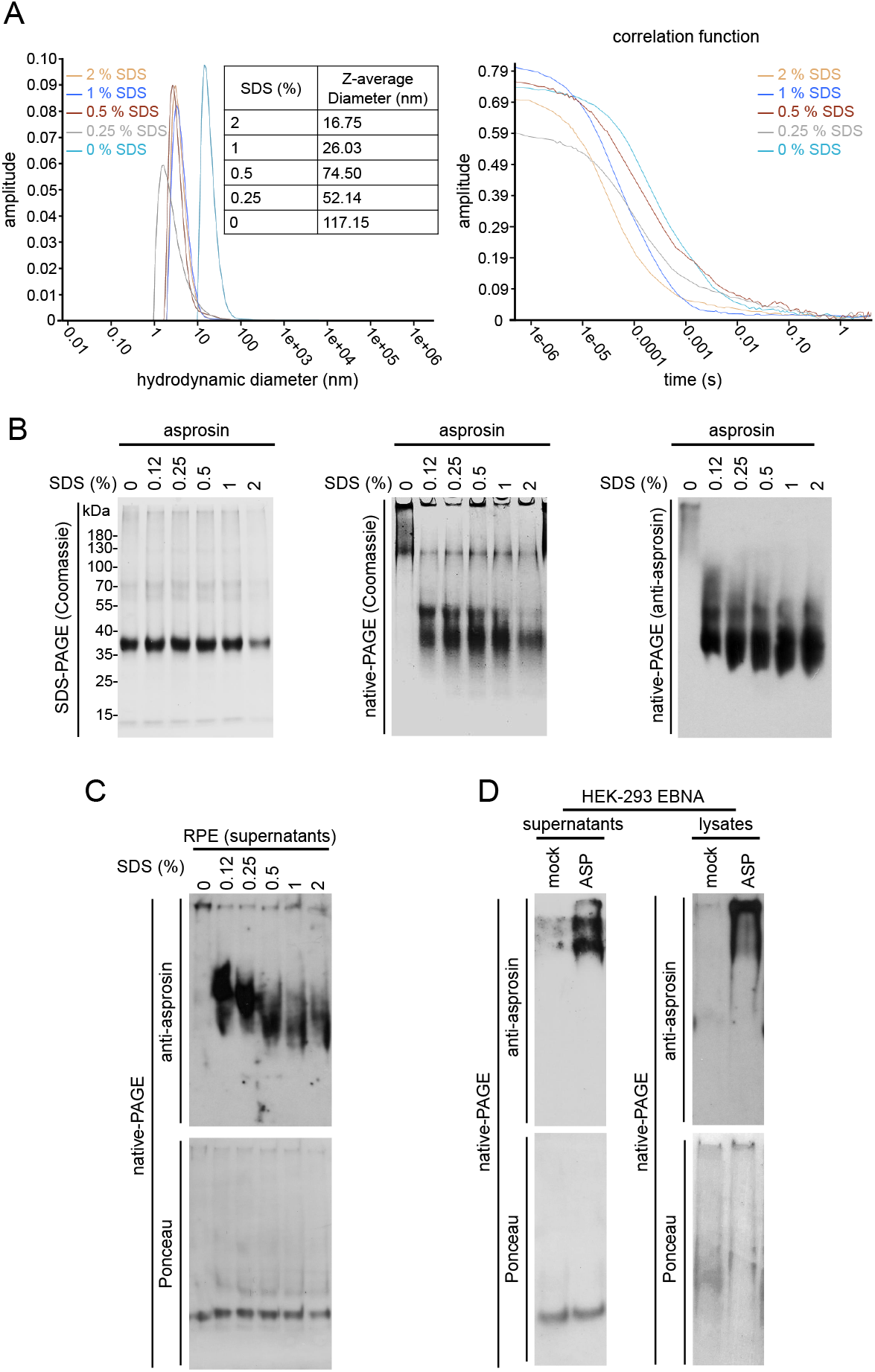
Asprosin forms multimers intracellularly. **A**. (left) Hydrodynamic diameters of asprosin in presence of 0-2% SDS determined by UNcle analyzer. Reduction of the hydrodynamic radius from 10-100 nm to 1-10 nm with varying concentrations of SDS reflects DLS measurements (Fig. 3C). (right) Correlation functions of asprosin in presence of 0-2% SDS by UNcle indicating dissociation of asprosin multimers. **B**. Treatment of recombinantly purified asprosin with serial concentrations of SDS. (left) Coomassie staining of SDS-PAGE gel after subjecting untreated and SDS-treated asprosin shows no change in asprosin band pattern after SDS treatment. (middle, right) Coomassie staining and western blot analysis of untreated and SDS-treated asprosin subjected to native-PAGE show dissociation of asprosin multimers. **C**. Treatment of RPE cell culture supernatants with 0-2% SDS followed by 10% native-PAGE and western blot analyses. SDS-treatment resulted in dissociation of endogenous asprosin multimers similar to SDS-treated recombinant asprosin (B, right). **D**. Western blot analysis of cell culture supernatants (left) and cell lysates (right) from stably transfected HEK-293 EBNA asprosin overexpressing cells. Supernatants and cell lysates of mock and asprosin transfected cells were subjected to 10% native-PAGE followed by western blot analysis. Immunoblots show accumulation of overexpressed asprosin at stacking gel indicating that asprosin multimerization already occurs intracellularly.

To examine whether asprosin oligomerization takes place intra- or extracellularly, cell-lysates and supernatants of RPE cells and stably transfected asprosin overexpressing HEK-293 cells were analyzed. Western blot analysis of cell culture supernatants of RPE cells after native-PAGE demonstrated the presence of endogenous asprosin in the stacking gel, which migrated into the gel after SDS treatment to the same positions as recombinant asprosin (Fig. 4C). Similarly, we observed the presence of asprosin multimers in the supernatant as well as cell lysates of asprosin overexpressing HEK293 cells by native SDS-PAGE (Fig. 4D), which dissociated upon SDS treatment (supplementary Fig. S5A).

### Asprosin is a substrate of tissue transglutaminase

Resistance of asprosin multimer bands to reducing agents (Fig. 2A, C) indicated stable covalent linkage between monomers. Fibrillin monomers are known to be crosslinked by transglutaminase 2 (TG2) which facilitates the assembly of stable fibrillin monomers (18). In general, transglutaminases primarily catalyze the formation of an isopeptide bond between γ-carboxamide groups of glutamine residue side chains and the ε-amino groups of lysine residue side chains with subsequent release of ammonia. Thereby, transglutaminases are very specific for the recognition of the glutamine residue and also depend on the charge of the flanking amino acid sequence (19). Since musculoskeletal cells are known to express transglutaminases (20), we tested whether asprosin is a substrate of TG2. Addition of TG2 to monomeric asprosin resulted in a significant shift of asprosin positive immunoblot signals towards higher molecular weight positions (Fig. 5A). A similar pattern in of bands representing asprosin oligomers was obtained when monomeric asprosin was incubated the presence of the chemical crosslinker disuccinimidylsuberat (DSS) (Fig. 5B). To confirm that asprosin is a substrate of TG2, recombinant asprosin and the co-substrate monodansylcadaverine (MDC) were incubated with TG2. Thereby, any TG2 substrate would become covalently tethered to MDC which allows sensitive detection by UV light (Fig. 5C). Subsequent to asprosin incubation with MDC and TG2, we were able to visualize monomeric and dimeric asprosin by UV light, thereby demonstrating that asprosin is a substrate of TG2 (Fig. 5D). Since covalent crosslinking of MDC adds 335.5 Da to the molecular weight of asprosin, we wanted to identify the glutamine residues via which asprosin crosslinking is mediated. Therefore, we subjected MDC tethered monomeric asprosin to a proteolytic digestion by trypsin and Lys-C and analyzed the resulting fragments by mass spectrometry analysis. Our analysis identified the glutamine residues Q99 and Q113 to be critical for TG2 mediated asprosin oligomerization (Fig. 5E).

**Figure 5:**
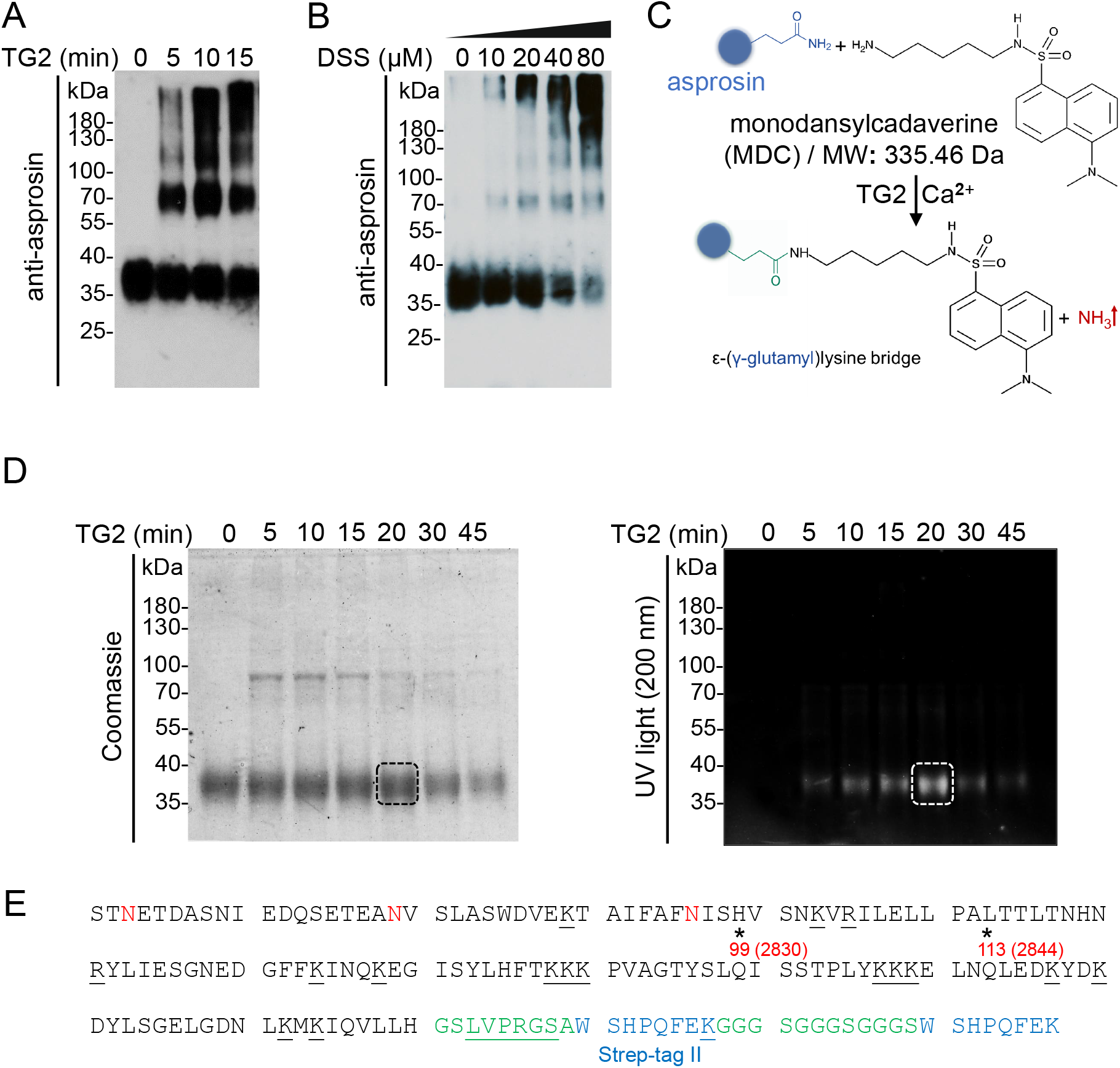
Asprosin is a substrate of transglutaminase 2 (TG2) **A**. Incubation of monomeric recombinant asprosin with TG2 results in a significant shift of asprosin positive immunoblot signals towards higher molecular weight positions. **B**. Incubation of monomeric recombinant asprosin with a chemical crosslinker, DSS shows the same higher molecular weight bands as after incubation with TG2. **C**. The crosslinking reaction between asprosin and fluorescent monodansylcadaverine (MDC), a TG2 substrate. **D**. Incubation of monomeric recombinant asprosin with monodansylcadaverine (MDC) in the presence of TG2 results in bands corresponding to monomeric and dimeric asprosin detected by UV light. The band marked by a dashed box was analyzed by mass spectrometry to identify the potential glutamine residues subjected to TG2 modification. **E**. Mass spectrometry analysis revealed the glutamine residues (black asterisks, marked in red, position regarding the fibrillin-1 sequence in parentheses) which could be utilized by TG2 for asprosin. Residues representing linker regions are indicated in green, thrombin cleavage (LVPRGS) site is underlined, and Strep-tag II sequences are marked in blue. The underlined residues mark predicted cleavage sites of Lys-C and trypsin used to generate peptide library of MDC-crosslinked asprosin. Anti-asprosin (pc-asp Ab) was used in the presented immunoblots in A and B.

### Asprosin fiber formation requires the presence of an intact fibrillin fiber network

Since our data demonstrated an intrinsic ability of asprosin to form multimers, we wanted to investigate whether asprosin fiber formation does also occur in the absence of cells (Fig. 6). For this purpose, we incubated asprosin in presence or absence of TG2 with or without SDS pre-treatment. We observed that asprosin forms fibers that were detectable by immunofluorescence in absence of TG2. However, TG2 addition significantly promoted the formation of an elaborate asprosin fiber network of considerable density and thickness. SDS pre-treatment blocked asprosin fiber formation in absence but not in presence of TG2, although the fibers appeared to be thinner and less dense (Fig. 6).

**Figure 6.**
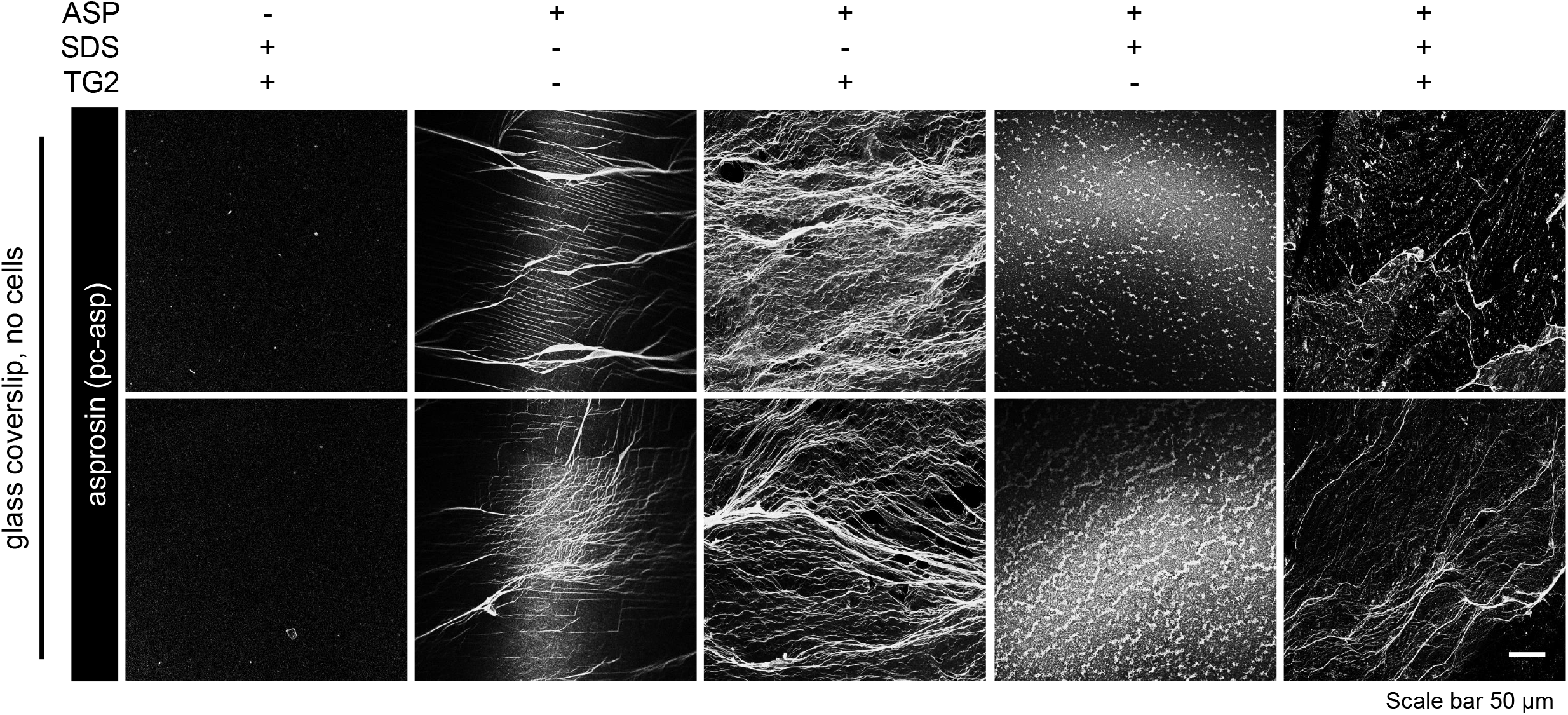
Asprosin fiber formation in absence of cells. Formation of asprosin fibers after incubating recombinant asprosin (8 μg/coverslip) for 48 h in presence or absence of TG2 (1μg/ml) with or without SDS (0.001%) pre-treatment. Asprosin fibers were detected after incubating with anti-asprosin (pc-asp Ab) (white). Images were obtained from a Leica SP8 confocal microscope and were processed using Leica Application Suite X (LAS X) software (version 3.7.5.2) and Fiji/ImageJ software (version 1.53t) to obtain average intensity Z-projection. Scale bars: 50 μm.

To understand whether asprosin fiber formation is blocked or promoted by the presence of extracellular scaffolds assembled by cells, we administered recombinant asprosin to cell cultures of several cell types showing high and low TG2 expression levels (Fig. 7A) followed by immunofluorescence analysis (Fig. 7B, C). While asprosin addition led to the formation of aggregates in cultures of U2OS cells, it resulted in robust formation of asprosin positive fibers in RPE cells or primary skin fibroblast cultures, while no fiber formation was detected in HepG2 or HEK-293 cells (Fig. 7C, supplementary Fig. S6A). Interestingly, asprosin positive fibers showed an overlapping distribution to the fibronectin and fibrillin-1 networks (Fig. 7C).

**Figure 7:**
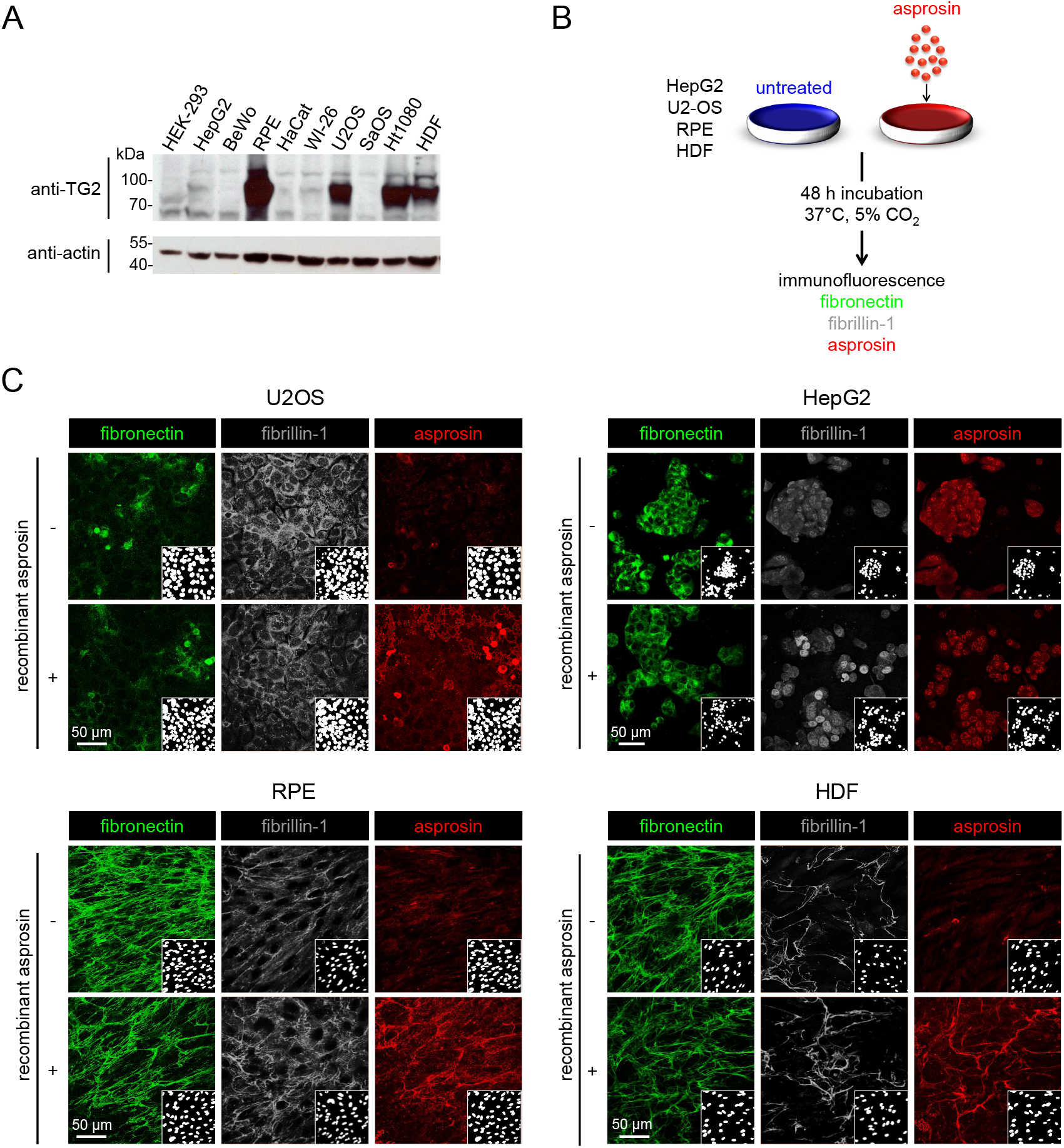
Assembly and ECM deposition of administered recombinant asprosin. **A**. (top) Western blot analysis showing the corresponding TG2 expression in cell lines lysates. (bottom) β□actin was used as a loading control. **B**. Schematic diagram illustrating the experimental design of cell lines treated with recombinant asprosin followed by immunofluorescence analysis. **C**. Recombinant asprosin (5μg/ml ∼ 140 nM) was administered to cell cultures of (top-left) U2OS, (top-right) HepG2, (bottom-left) RPE, and (bottom-right) HDF cells. 48 h after asprosin treatment, cells were immunostained with anti-fibronectin (FN1 Ab) (green), anti-fibrillin-1 (FBN1, rF90 Ab) (gray), anti-asprosin (pc-asp Ab) (red), and DAPI (white, nuclei). Immunofluorescence analysis reveals the presence of intact fibrillin-1 and fibronectin fiber network in cultures of RPE and HDF cells, but not in cultures of U2OS and HepG2 cells. Asprosin positive fibers were observed after administration to RPE cells or HDF. However, asprosin is detected as aggregates in asprosin-treated U2OS cells, while HepG2 shows no significant differences in staining patterns before and after asprosin treatment. Images were obtained from a Leica SP8 confocal microscope and were processed using Leica Application Suite X (LAS X) software (version 3.7.5.2) and Fiji/ImageJ software (version 1.53t) to obtain average intensity Z-projection. Scale bars: 50 μm.

Since we previously determined that about 20 kDa of the 37 kDa monomeric asprosin is due to glycosylation (14), we wanted to investigate whether glycosylation is required for asprosin fiber formation. However, immunofluorescence analysis showed that de-glycosylated asprosin is fully capable to form fibers in RPE cell cultures (supplementary Fig. S6B, C), suggesting a requirement for the core protein only. To address the question whether non-covalent multimerization is required for asprosin fiber formation, we incubated asprosin with SDS prior to the addition to cultures. However, our analysis showed that asprosin fiber formation was not affected by SDS pretreatment indicating that non-covalent association into multimers is not necessary for asprosin fiber formation (supplementary Fig. S6B, C). However, we did not observe fiber formation when placensin was added to fibroblast cultures (supplementary Fig. S3D).

Since TG2 is known to establish functional crosslinks thereby stabilizing fibronectin and fibrillin microfibril networks (21,22), we wanted to test whether TG2 may also tether asprosin to either of them. Pull down experiments revealed that asprosin interacts with the N-terminal half of fibrillin-1 and not with the C-terminal half of fibrillin-1 or full-length fibronectin (Fig. 8A, B). Also, interaction studies employing surface plasmon resonance (SPR) confirmed that asprosin binds to the N-terminal region of fibrillin-1 and not to full length fibronectin (Fig. 8C). To test whether intact fibrillin fiber assembly is required for asprosin fiber formation, we added recombinant asprosin to primary skin fibroblast cultures of MFS patients which show a compromised fibrillin fiber formation but a normal fibronectin assembly. Immunofluorescence analysis showed that asprosin fiber formation was abolished when fibrillin fiber formation was deficient (Fig. 9). To demonstrate the role of TG2 in asprosin fiber formation, we generated a mutant form of asprosin in which the identified crucial residues for transglutaminase-mediated crosslinking Q99 and Q113 were mutated to alanine (Fig. 10A). Upon addition of mutant asprosin to RPE cells, we found a significant reduction in fiber formation compared to the control culture incubated with wild-type asprosin (Fig. 10B), suggesting that Q99 and Q113 are crucial for fiber formation. To investigate whether asprosin is also targeted to the fibrillin-1 fiber network *in vivo*, we co-labelled tissue sections for fibrillin-1 and asprosin. By employing newly generated antibodies specific for the detection of endogenous asprosin in human and murine tissues employing, we could show that asprosin co-localizes with fibrillin fibers in connective tissues such as skeletal muscle, heart, and cruciate ligament (Fig. 11).

**Figure 8:**
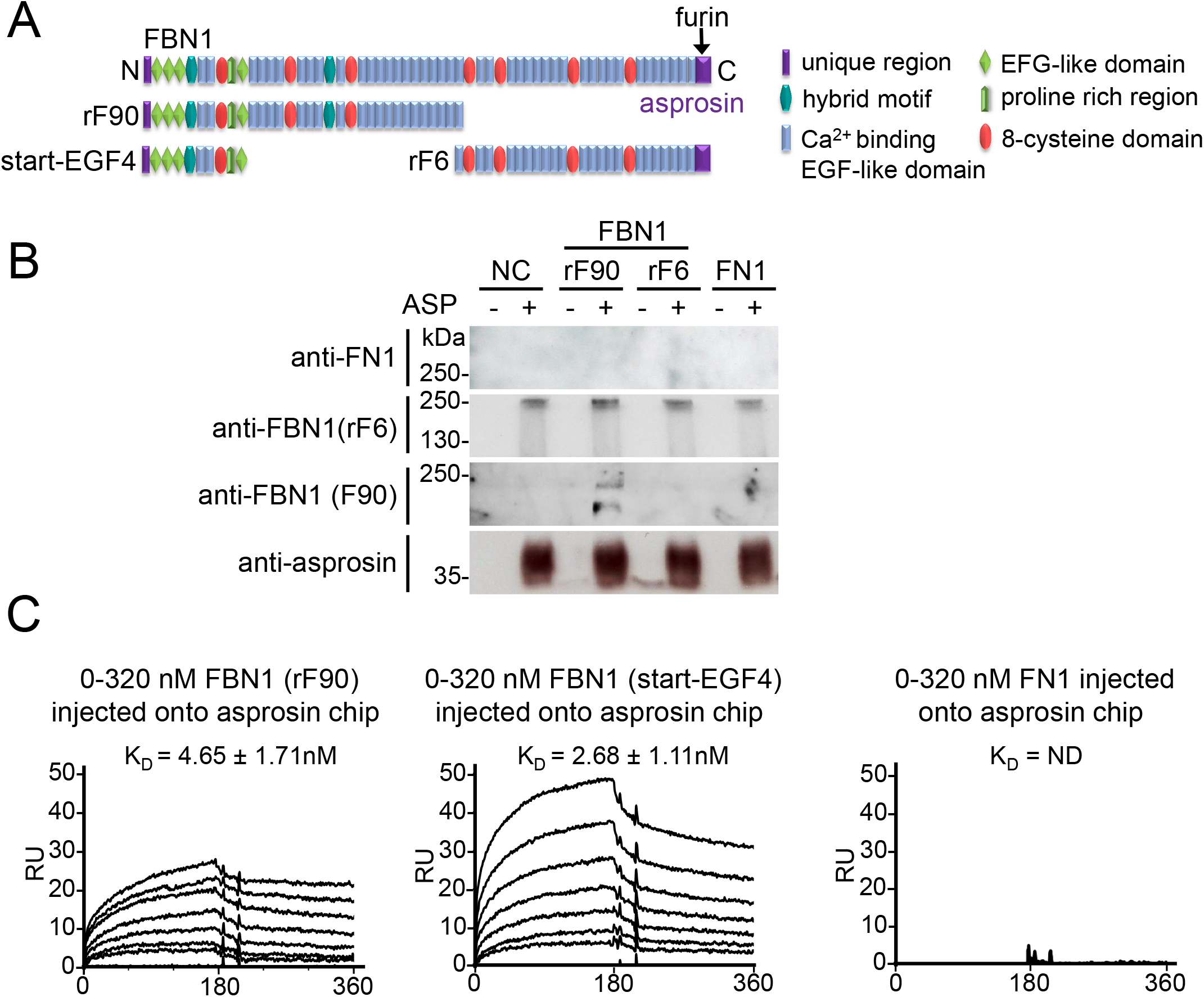
Asprosin interacts with the N-terminal region of fibrillin-1. Domain structure of fibrillin-1 and its N- and C-terminal halves including the start-EGF4 fragment (furin cleavage site marked by arrow). **B**. HEK-293 EBNA cells were transfected with overexpression constructs encoding for fibrillin-1 N- and C-terminal halves (rF90 and rF6). Medium obtained from non-transfected HEK-293 EBNA cells served as negative control (NC). Media obtained from the transfected with rF90 and rF6 are indicated. HEK-293 EBNA (NC) medium supplemented with fibronectin (10 μg) is indicated as FN1. The media were subjected to incubation with recombinant asprosin (double C-terminal 2×Strep-tag II) in presence of 0.5 μg/ml TG2 overnight at 4°C. The mixtures were then subjected to pulldown with Strep-Tactin XT beads. After washing, the eluted fractions were subjected to immunoblot analysis with the indicated antibodies revealing the binding of asprosin to N-terminal half of fibrillin-1 (rF90). **C**. SPR binding studies suggest that asprosin interacts with the N-terminal half of fibrillin-1 (rF90) and the start-EGF4 fragment but does not bind to fibronectin.

**Figure 9:**
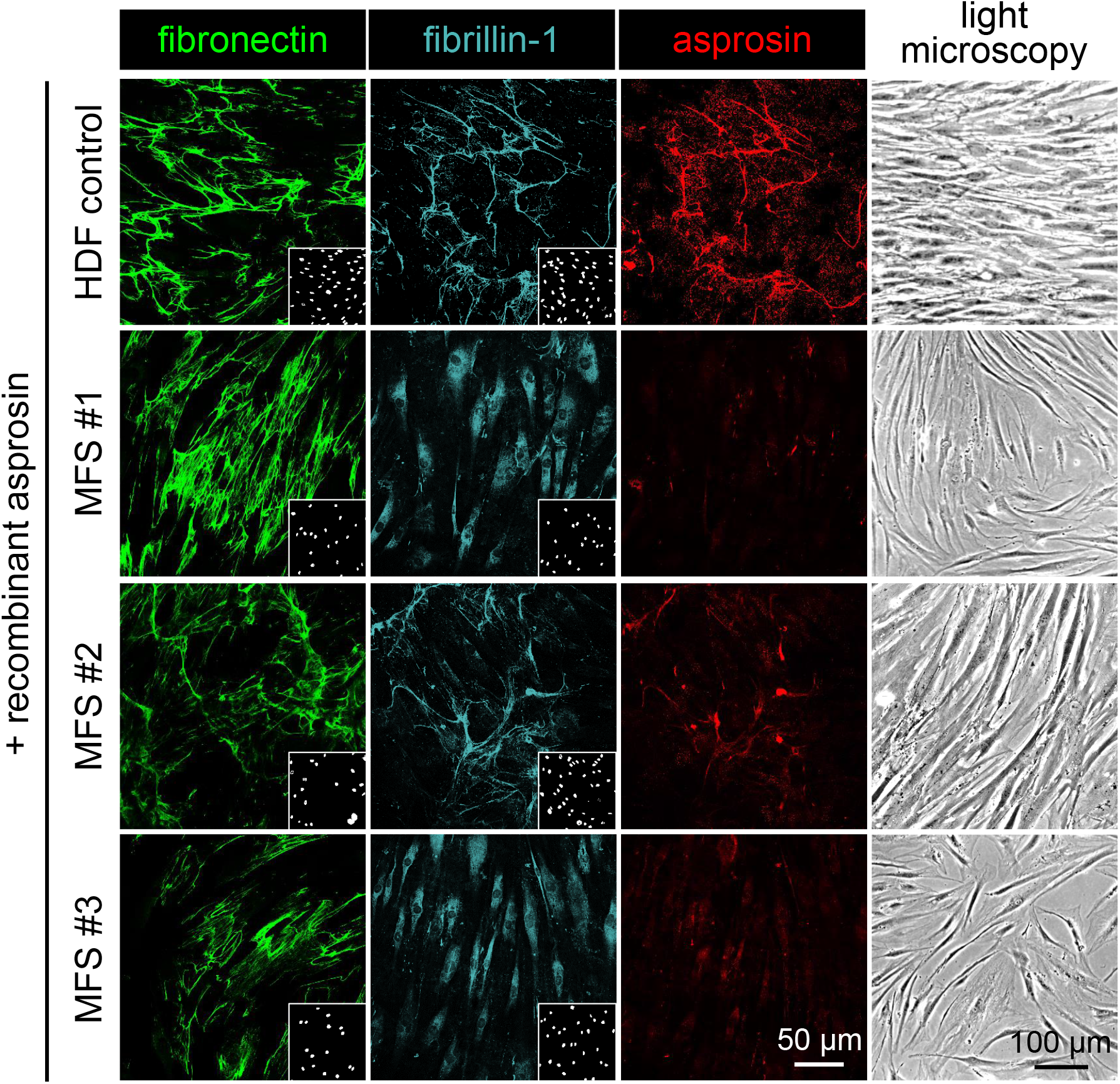
Linear deposition of asprosin as fibers is fibrillin-1 dependent. The formation of asprosin positive fibers after administration of recombinant asprosin (5μg/ml ∼140 nM) depends on the integrity of the fibrillin-1 fibril network which is deficient in primary fibroblast cultures from MFS patients. Control and MFS patient cells were incubated with anti-fibronectin (FN1 Ab) (green), anti-fibrillin-1 (FBN1, rF90 Ab) (gray), anti-asprosin (hASP(rat) Ab) (red), and DAPI (white, nuclei). Light microscopy images were obtained by Axiophot Microscope (Carl Zeiss, Germany). Fluorescence microscopy images were obtained from a Leica SP8 confocal microscope and were processed using Leica Application Suite X (LAS X) software (version 3.7.5.2) and Fiji/ImageJ software (version 1.53t) to obtain average intensity Z-projection.

**Figure 10:**
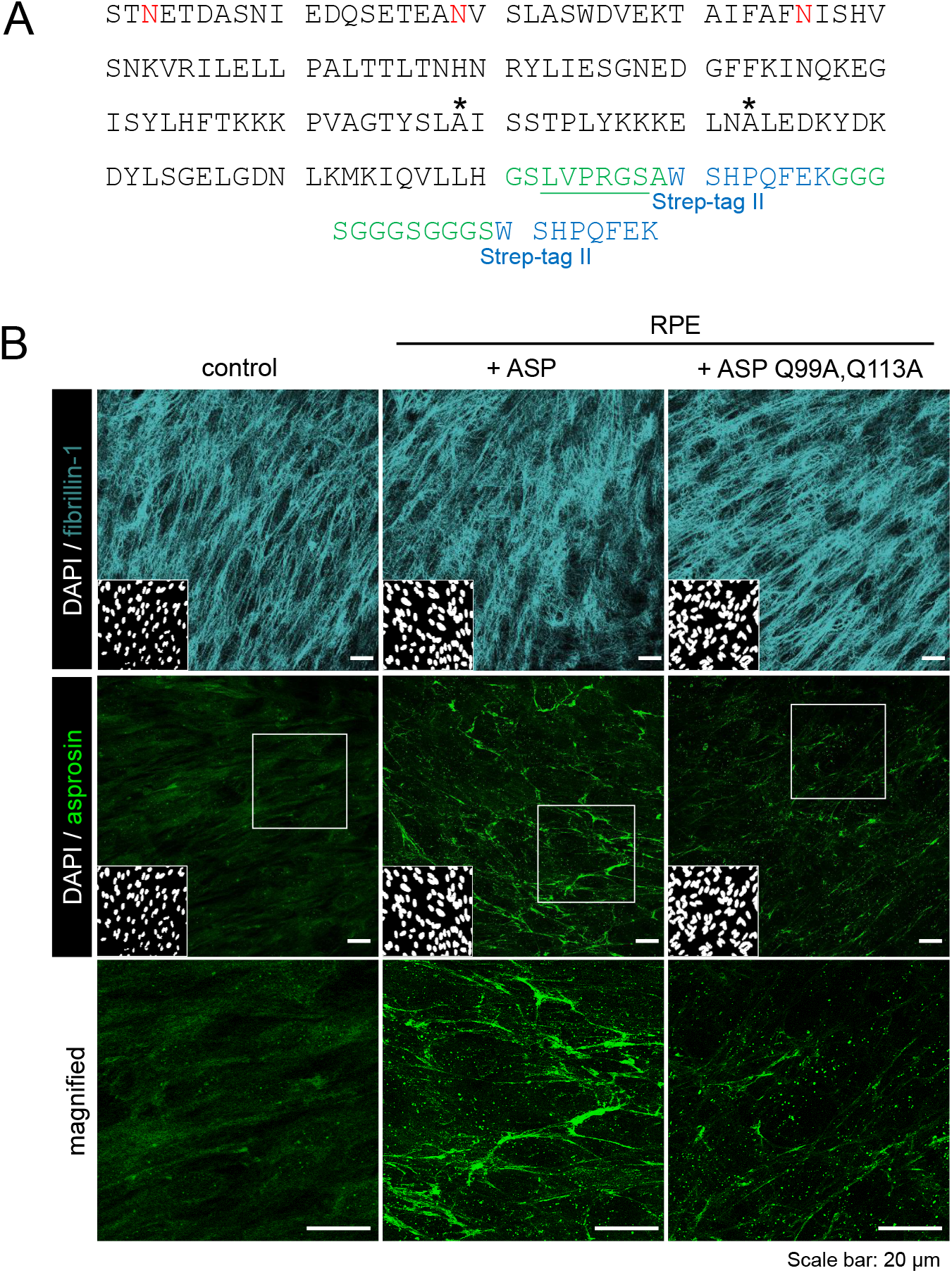
Mutation of critical glutamine residues abolishes asprosin fiber formation. **A**. Protein sequence of asprosin carrying glutamine substations (Q99A, Q113A, black asterisks). Residues representing linker regions are indicated in green, thrombin cleavage (LVPRGS) site is underlined, and Strep-tag II sequences are marked in blue. **B**. Comparative immunofluorescence analysis after administration of recombinant wild-type and mutated asprosin (5μg/ml ∼140 nM) to RPE cell culture. Untreated and treated RPE cells were incubated with DAPI (white, nuclei), anti-asprosin (pc-asp Ab) (green), and anti-fibrillin-1 (rF90 Ab) (gray). 3-fold magnified areas are marked by white boxes. Fibrillin-1 immunostaining (gray) reveals an intact fibrillin-1 network in untreated and treated RPE cells. (top panel) The immunofluorescence staining of asprosin in untreated cells (control) does not show asprosin positive fibers. However, (middle panel) the wild-type asprosin treated RPE cells show intact asprosin positive fibers compared to (bottom panel) mutated asprosin treated RPE cells, which show significantly disrupted fibers. Fluorescence microscopy images were obtained from a Leica SP8 confocal microscope and were processed using Leica Application Suite X (LAS X) software (version 3.7.5.2) and Fiji/ImageJ software (version 1.53t) was used to obtain average intensity Z-projection.

**Figure 11:**
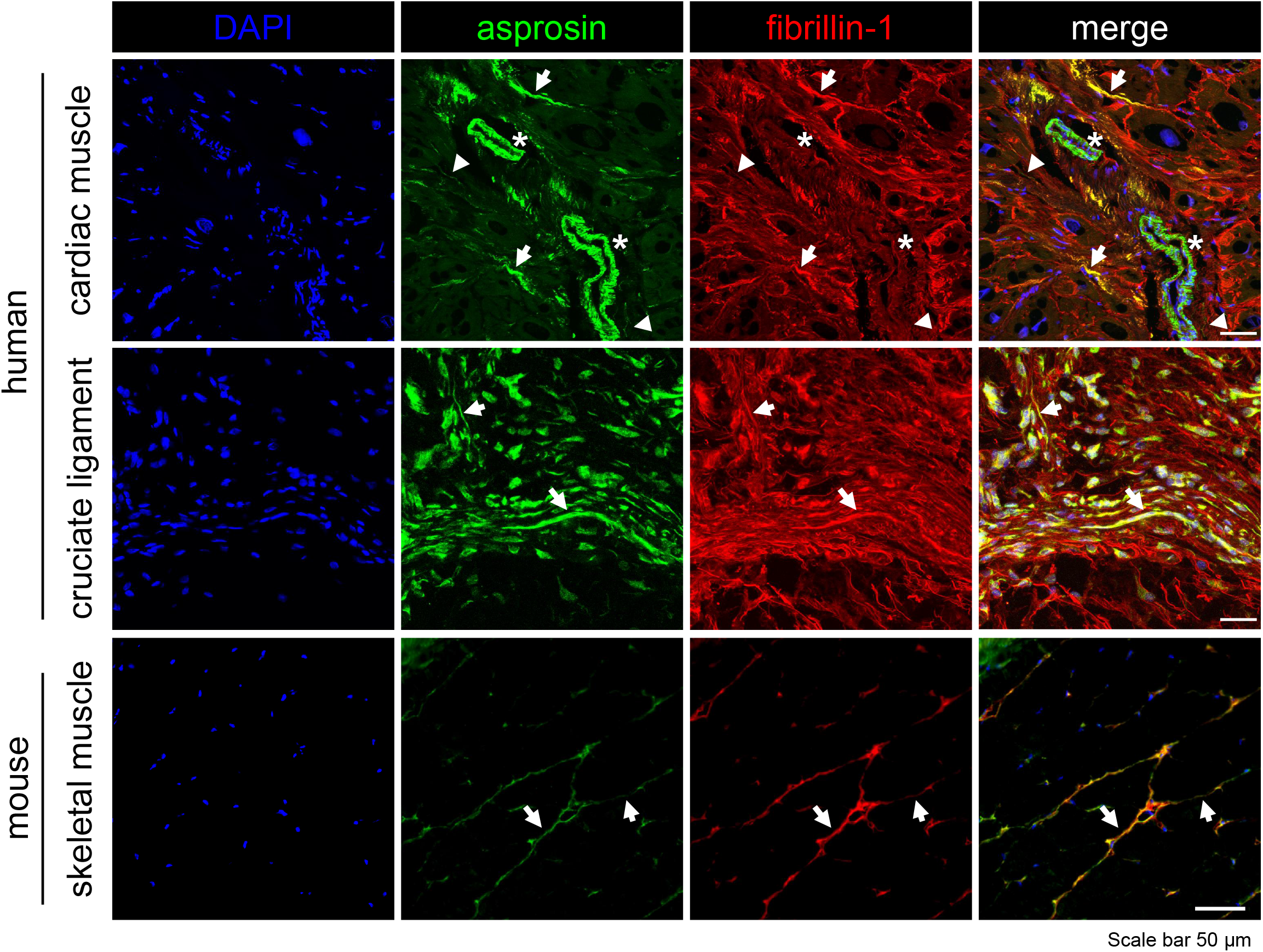
Asprosin colocalizes with fibrillin-1 in human and murine tissues. Confocal immunofluorescence microscopy of asprosin in indicated human and mouse tissues. Sections were fixed with acetone and incubated with asprosin antibodies (green; pc-asp Ab for human tissues, mASP(rat) Ab for murine tissues), anti-fibrillin (FBN1, rF90 Ab) (red) and DAPI (blue, nuclei). Immunostaining reveals depositions of asprosin positive fibers co-localizing with fibrillin-1 positive fibers (see merge panel). In cardiac muscle asprosin also appears as separate fine structures (arrowhead). Marked immunoreactivity for asprosin is present in the cytoplasm of vascular smooth muscle cells (VSMCs) and fibroblasts (asterisks). Images were obtained from an Olympus BX43 microscope (Olympus) equipped with a DP80 dual CCD camera and Leica SP8 confocal microscope. Images were processed using Leica Application Suite X (LAS X) software (version 3.7.5.2) and Fiji/ImageJ software (version 1.53t) was used to obtain average intensity Z-projection.

### Asprosin multimerization affects cellular uptake

To examine whether asprosin multimerization affects its internalization, we incubated several cell types including HEK-293, WI-26, and RPE with asprosin, in presence and absence of SDS for one hour followed by western blot analysis. Our analyses demonstrated that SDS mediated dissociation of non-covalently associated asprosin multimers significantly promoted asprosin uptake into RPE, WI-26, and HEK-293 cells (Fig. 12A, B). Interestingly, in the analyzed cell lysates we could not only detect monomeric asprosin at ∼37 kDa but also bands corresponding to asprosin multimers (e.g. at ∼70 kDa corresponding to an asprosin dimer) (Fig. 12B, C). This finding indicated a potential multimerization of asprosin during the uptake process, since asprosin multimers were only detectable in minimal amounts prior to incubation with the cell layer (Fig. 12).

**Figure 12:**
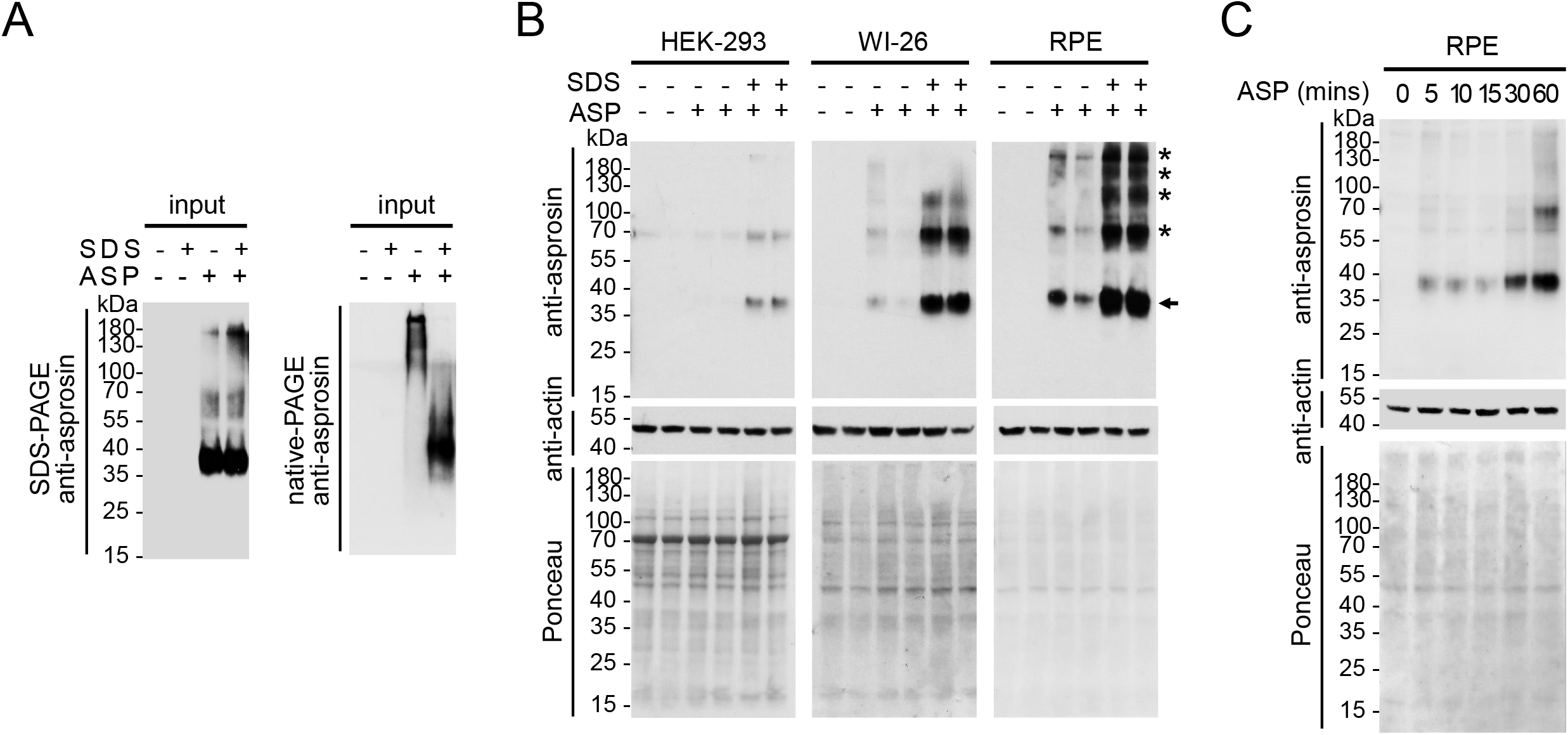
Asprosin oligomerization affects its cellular uptake. **A**. Quality control immunoblots of recombinant asprosin protein preincubated with SDS and diluted into cell culture media was subjected to (left) 10% SDS-PAGE gel and (right) 10% native-PAGE gel. Dissociation of asprosin multimers after SDS treatment can be only observed in samples subjected to native-PAGE. **B**. Comparative western blot analysis with anti-asprosin (pc-asp Ab) of cell lysates collected from (left) HEK-293, (middle) WI-26, and (right) RPE cells after administration of recombinant asprosin (3.5 μg/ml∼100 nM) in presence and absence of SDS. Immunoblots show the enhancement of monomeric asprosin uptake in after SDS in all treated cell lines (band ∼37 kDa, black arrow), additionally, in WI-26 and RPE cells asprosin positive bands of higher molecular weight (black asterisks) were also detected suggesting multimerization of administrated asprosin. **C**. Immunoblot with anti-asprosin (pc-asp Ab) of cell lysates collected from RPE cells treated with recombinant asprosin (1.75 μg/ml ∼ 50 nM) in presence of SDS after certain time points (5, 10, 15, 30, and 60 min). A significant increase of monomeric asprosin signal (band ∼37 kDa) within time was observed and asprosin positive band at 70 kDa was detected corresponding to dimeric asprosin signal. β-actin and Ponceau staining were used as loading controls.

### Molecular modelling of asprosin suggests a cadherin like fold

By employing computational approaches we were able to generate a structural model of asprosin by employing AlphaFold2 and ColabFold (23). The predicted asprosin model showed a high local structure confidence in the last 120 C-terminal residues (S^21^-H^140^) (Fig. 13A). In addition, we submitted the generated structure to Dali server, a protein structure comparison server (24) in order to identify structures similar to asprosin predicted structure in Protein Data Bank (PDB). The obtained data suggested that asprosin is predicted to have a cadherin-like fold (Fig. 13A).

**Figure 13:**
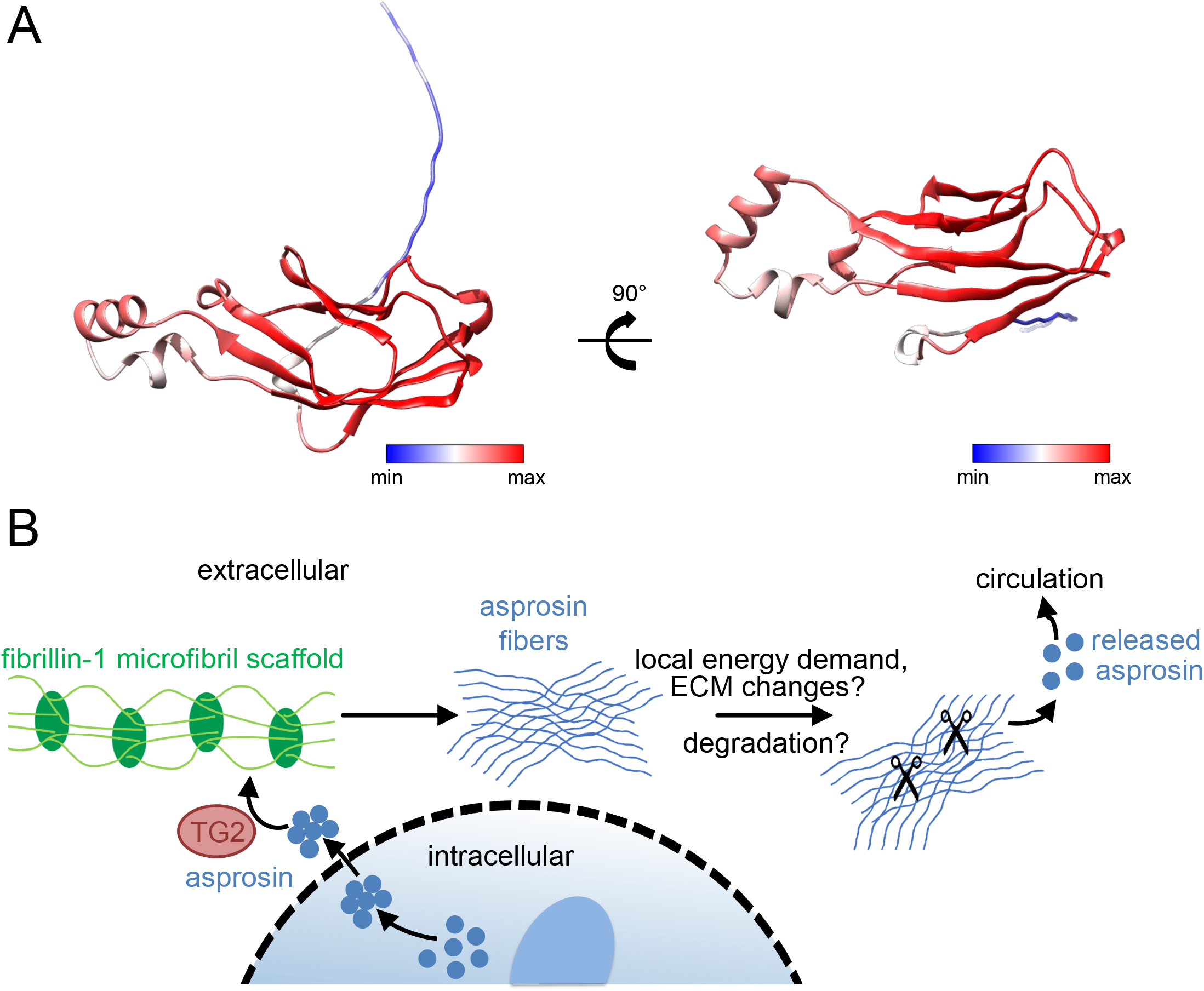
Mechanistic model for targeting of asprosin to the extracellular matrix. **A**. Model of asprosin. Asprosin assumes a cadherin-like core structure with the potential capability to multimerize via cadherin-like domains. The model is colored according to the predicted local distance difference test (pLDDT) and the per-residue confidence (red: high values, blue: low values). Image was rendered by employing ChimeraX v1.4 using the predicted model downloaded from Colab Alphafold2. **B**. The model shows a potential mechanism of asprosin storage and utilization within the extracellular matrix. Upon secretion of asprosin may be targeted to assembled fibrillin-1 fibers. Thereby, TG2 mediated crosslinking enables stable storage of asprosin in the form of multimers. Multimeric asprosin may be later utilized by specific activation mechanisms such as proteolytic degradation of asprosin positive fibers. Released asprosin may act locally or is transported via the circulation to other target organs.

## Discussion

Currently, asprosin is mainly viewed as WAT derived hormone that is released into the blood stream to regulate glucose release in the liver (7,13). However, our findings suggest a new mechanism of asprosin storage and utilization within the ECM (Fig. 13 B). Once secreted, asprosin is targeted to assembled fibrillin fibers, whereby TG2 mediated crosslinking enables stable storage of asprosin in the form of multimers. Multimeric asprosin may be later utilized by specific activation mechanisms such as proteolytic degradation of asprosin positive fibers. Released asprosin may act locally or is transported via the circulation to other target organs.

However, since asprosin is the C-terminal propeptide of profibrillin we wondered whether it has a similar tissue distribution as fibrillin-1. By generating new asprosin specific antibodies that do not crossreact with full length fibrillin-1 (14) we were able to detect extracellular deposits of asprosin as fibers in tissues. This finding is in line with previously reported mass spectrometry data from extracted mature microfibrils isolated from the human zonular apparatus (Cain et al. 2006). In this preparation two verified fingerprint peptides of asprosin downstream of the C-terminal furin cleavage site sequence were reliably detected (Cain et al., 2006). This finding was unexpected since profibrillin-1 processing by furin is demonstrated as a prerequisite for mature fibrillin-1 deposition (25-27). Asprosin positive fibers were more prominently observed in certain tissue microenvironments such as skeletal or cardiac muscle. However, in liver, asprosin was mainly found in intracellular vesicles (Fig. 1) indicating its rapid utilization in this metabolically active tissue.

At present, there is limited structural data on asprosin. Here, we propose a structural model by using AlphaFold suggesting a conformation similar to extracellular cadherin (EC) repeats. Cadherins are known to facilitate self-oligomerization to ensure tissue cohesion. Thereby, cadherins are known to mediate calcium-dependent cell-cell adhesion by forming trans-interactions via EC1 domains from to each other attached cells (28,29). In addition, a cis-interface involving the nonsymmetrical interaction of the EC1 domain and the EC2 domain of neighboring cadherins was proposed (30). The structural similarity of asprosin to EC repeats may endow asprosin with the capability to align into multimers in a similar fashion as cadherins (Fig. 13A). The formation of such stable multimers may then be required for further stabilization via covalent bonds by TG2 (Fig. 9A). Chelation of extracellular Ca^2+^ is known to inhibit the adhesive binding activity of cadherin ectodomains and thereby disrupts epithelial cohesion (31). However, our attempts to dissolve asprosin multimers by complexing calcium via EDTA treatment were not successful (supplementary Fig. S5F), suggesting that the presence of Ca^2+^ does not contribute to the stability of asprosin multimers.

So far, the function of asprosin fibers is unknown. We hypothesize that the oligomerization and linear deposition of asprosin as fibers represents a local asprosin storage mechanism within tissue-specific microenvironments (Fig. 9B). Asprosin stored in tissues may function as a sensor for local energy demand. From there it may be released by specific, yet unknown activation reactions into the serum. Controlled asprosin release by tissues may be a so far unexplored mechanism how tissues signal energy demand to the metabolic system. A metabolic function of asprosin in the regulation of the energy demand of tissue resident cells was already proposed in *in vitro* cell cultures. For instance, experiments with murine myoblasts (C2C12) showed that asprosin is not only able to interfere with muscle cell insulin sensitivity (12), but also up-regulates glucose transporter 4 (GLUT4) expression in C2C12 myotubes and thereby enhance local glucose uptake by tissue muscle cells (32).

We found that asprosin is a specific substrate of TG2, a crosslinking enzyme that is known to facilitate fibronectin, fibrillin-1, and elastic fiber assembly (18,21,33,34). TG2 is very selective regarding the glutamine residues it subjects to crosslinking (19). Our experiments showed that placensin, the C-terminal propeptide of fibrillin-2, does not form higher oligomers or fibers in fibroblast cultures upon addition, despite the presence of several available glutamine residues (supplementary Fig. S3D). Recently, placensin was described as a new placenta-derived glucogenic hormone that stimulates hepatic cAMP production, protein kinase A (PKA) activity and glucose secretion (16). Also asprosin was shown to be expressed in human placenta and is elevated in the plasma of pregnant women complicated with gestational diabetes mellitus (GDM) and their offspring (umbilical blood) (35) after adjustment for maternal and neonatal clinical characteristics and lipid profiles. In placenta, asprosin fiber formation may have the function to fine tune the metabolic effects mediated by asprosin and placencin.

Our transfection experiments with HEK293 cells constructs encoding monomeric asprosin showed that asprosin multimers were also detected intracellularly, whereas in the conditioned media only monomeric asprosin was detected. In contrast, in the lysates and supernatant of 3T3-L1 cells, asprosin oligomers were found intra- and extracellularly. Intracellular TG cross-linking is a complex theme as TGs do not pass through the conventional ER/Golgi route and require Ca^2+^ for activity. Nevertheless, despite low intracellular calcium levels, multiple transamination and crosslinking substrates of intracellular TG2 have been identified (20,36). This suggests that locally increased intracellular calcium and/or as yet uncharacterized interacting proteins may facilitate formation of active TG2. Our transfection studies with an overexpression construct coding for monomeric asprosin already indicated the formation of tetramers in cell lysates (Fig. 2B). However, native-PAGE analysis of cell lysates from asprosin overexpressing HEK293 cells indicated that non-covalently associated multimers already form intracellularly (Fig. 4D). This suggests that the intrinsic ability of asprosin monomers to form very stable multimers allows the establishment of covalent crosslinks via TG2 within the intracellular milieu despite low calcium concentrations. Also, internalization studies using monomeric asprosin that was pre-treated with SDS suggest asprosin undergoes multimerization upon uptake to the intracellular space. Uptake of monomeric asprosin was more sufficient, but multimer bands of intracellular asprosin pools were prominent subsequently (Fig. 12). These results indicate that asprosin multimerization does limit cellular uptake of asprosin likely due to reduced access of relevant epitopes recognized by cell surface receptors. It has been previously reported that protein multimerization can affect its cellular uptake and regulate pathogenesis of several diseases (37-40). For example, multimerization of Tau proteins and α-Synuclein is involved in the pathogenesis of neurodegenerative diseases such as Alzheimer’s and Parkinson’s diseases (41-44).

In future studies we will address not only further investigate the molecular requirements for cellular utilization of asprosin cells, but also by what specific release mechanisms asprosin may be liberated from its fibrous state.

Our data show that asprosin is specifically targeted to fibrillin-1 fibers and not to fibronectin. This finding may implicate an intact fibrillin network in the proper extracellular storage of asprosin. Fibrillin-1 is ubiquitously expressed in all tissues and is known to assemble into supramolecular microfibrils that serve as targeting scaffolds for connective tissue derived growth factors such as TGF-β and BMPs (45-49). By targeting and sequestration of asprosin in connective tissue microenvironments fibrillin-1 may be involved in the regulation of spatio-temporal energy demand of tissue resident cells.

## Materials and method

### Ethics statement

The use of human specimen involved in this study was approved by the institutional review boards at the Medical Faculty of the University of Cologne, the German Sports University, and Ghent University. Written informed consent was obtained from patients in accordance with institutional review board policies. Written informed consent was obtained from all participating probands and patients who agreed to have materials examined for research purposes. The study was conducted in accordance with the Declaration of Helsinki. Human samples were collected according to the Biomasota code (13-091) at the University Hospital of Cologne, Cologne, Germany and Ethics Committee of the Medical School Brandenburg, Brandenburg, Germany (reference numbers: E-01-20200921, E-01-20211115). The study was approved by the ethics committee of the University Hospital of Cologne (18-052). Mice were sacrificed by cervical dislocation and immediately dissected for experiments. All animal procedures were conducted in compliance with protocols approved by the Committee on the Ethics of Animal Experiments of the Landesamt für Natur, Umwelt und Verbraucherschutz Nordrhein-Westfalen (84-02.04.2019.A326) and were in accordance with National Institutes of Health guidelines.

### Antibodies

Pc-asp anti-asprosin, fibrillin-1 (rF90) polyclonal antibody, and fibrillin-1 rabbit monoclonal antibody (CPTC-FBN1-3) (DSHB, Iowa, USA) were previously described (14). Monoclonal anti-fibronectin antibody (# F7387) was purchased from Merck Millipore (Massachusetts, USA). Mab anti-asprosin antibody (clone Birdy-1, AG-20B-0073) was purchased from AdipoGen Life Sciences Inc. (San Diego, USA). StrepMAB-Classic antibody (#2-1507-001) was from IBA GmbH, Germany. Antibodies against murine asprosin (mASP(rb) Ab, (mASP(rat) Ab) were generated against full-length mouse asprosin (supplementary Fig. S3). Recombinantly expressed mouse asprosin or human placensin were used to immunize rabbits for polyclonal antibody production (Davids Biotechnologie GmbH, Regensburg, Germany). Recombinantly expressed human asprosin (14) was used for polyclonal antibody production in a rat (Pineda, Berlin, Germany). Preimmune serum samples (0.5 ml) were obtained before the immunization and did not show cross-reactivity to asprosin tested by ELISA. Antisera were affinity purified on CNBr-activated Sepharose 4B (Cytiva, Uppsala, Sweden) columns conjugated with the respective recombinantly expressed proteins (300 μg) according to the manufacturer’s instructions. Antibodies were eluted with 0.1 M glycine (pH 2.5) and neutralized with 3 M Tris/ HCl, pH 8. Eluted antibodies were concentrated by using Amicon Ultra Centrifugal Filters (cut-off: 10 kDa), (starting volume: 6 ml, end volume: 1ml). The specificity of the raised antibodies was tested by direct ELISA, pull down experiments, as well as immunofluorescence of tissues and transfected cells (supplementary Fig. S2-S5).

### Expression and purification of recombinant proteins

Human asprosin, mouse asprosin, human placensin, mutant asprosin (Q99A and Q113A), and the N-terminal half of fibrillin-1 (rF90, amino acids positions: (M^1^-V^1527^) were produced and purified as previously described (14) (supplementary Fig. S2B). C-terminal half of fibrillin-1 (rF6, amino acid positions V^1487^-H^2871^) was overexpressed in HEK-293 EBNA cells. The rF6 overexpression construct (50) was a kind gift from Lynn Sakai, Oregon Health and Science University, Portland, OR, USA. cDNAs encoding for mouse asprosin, human placensin, and mutant asprosin (Q99A and Q113A) were generated by GeneArt Strings DNA Fragments service (Thermo Fisher Scientific, Massachusetts, USA). including the restriction enzymes sequences for ends. Subcloning of cDNA sequences via NheI/ XhoI restriction sites into a modified pCEP-Pu vector as well as transfection of overexpression constructs in HEK293 EBNA cells followed by establishment of stably transfected cell clones, and protein purification from conditioned media via affinity chromatography using a C-terminally placed 2×Strep-tag II were as previously described (14).

### Size exclusion chromatography with multi-angle static light scattering (SEC-MALS) and electron microscopy (EM)

Recombinant asprosin (0.5 ml, 1 μg/μl) was subjected to gel filtration using a Superose 6 Increase 10/300 GL column in PBS at 0.6 ml/min. The eluate was passed through a Wyatt DAWN Heleos II EOS 18-angle laser photometer with a Wyatt QELS detector for the measurement of hydrodynamic radius. Data were analyzed using Astra 6.1 (Wyatt, Santa Barbara, USA). Elution fractions from SEC-MALS were analyzed by EM as previously described (51).

### Immunoblotting

Cells were lysed in RIPA buffer (50 mM Tris HCl, 150 mM NaCl, 1.0% (v/v) NP-40, 0.5% (w/v) sodium deoxycholate, 1.0 mM EDTA, 0.1% (w/v) SDS and 0.01% (w/v) at a pH of 7.4) supplemented with cOmplete protease inhibitor cocktail (#11697498001, Sigma-Aldrich, Darmstadt, Germany) and PhosSTOP phosphatase inhibitors (#4906837001, Sigma-Aldrich, Darmstadt, Germany). Protein concentrations were measured using Pierce BCA Protein Assay Kit (#23225, Thermo Fisher Scientific, Massachusetts, USA). To harvest cell culture supernatants, cell layers were incubated with serum free medium when cells reached 90% confluency and subsequently collected after 48 h. The collected serum free medium was filtered and concentrated with Amicon Ultra Centrifugal Filters (cut off: 3 kDa), (starting volume: 2 ml, end volume: 50 μl). Cell lysates and cell culture supernatant samples analyzed by 7.5% or 10% SDS-PAGE under reducing (with β-mercaptoethanol) or non-reducing conditions, followed by transfer to 0.45 μm PVDF transfer membranes (Thermo Fisher Scientific, Massachusetts, United States). Membranes were blocked with Pierce Protein-Free (TBS) Blocking Buffer (#37585, Thermo Fisher Scientific, Massachusetts, United States) for one hour, then incubated with the appropriate dilutions of primary antibodies overnight at 4°C and subsequently incubated with secondary antibody, mouse anti-rabbit IgG-HRP conjugate or goat anti-mouse IgG-HRP conjugate, 1:5000 for 1 h. All antibodies were diluted into blocking buffer. Signals were developed with SuperSignal West Pico PLUS Chemiluminescent Substrate (#34579, Thermo Fisher Scientific, Massachusetts, USA).

Samples used in native-PAGE analysis were separated under non-denaturing conditions. Cells used for native-PAGE analysis were lysed using mechanical lysis (manual grinding) in non-denaturing (detergent-free) buffer (100 mM Tris/HCl pH 8.0 150 mM NaCl) supplemented with cOmplete protease inhibitor cocktail and PhosSTOP phosphatase inhibitors. Cell lysates and cell culture supernatants were mixed with native-PAGE loading buffer (62.5 mM Tris-HCl, pH 6.8, 40% glycerol, 0.01% bromophenol blue) without heating prior to native-PAGE.

### Cellular uptake experiments

Cells were seeded in 6-well plates until reaching 70% confluency and were kept in DMEM (serum free medium) for 24 h. 6-well plates were placed on ice for 20 minutes prior to treatment with asprosin (50 or 100 nM) that was preincubated with or without SDS. For SDS treatment, 9 μl asprosin stock solution were mixed with 1 μl 1% SDS, which was further diluted into cell culture media to reach the indicated final asprosin concentrations. Thereby the final SDS concentration was 0.001% for which no cell toxicity was observed. After incubation with asprosin, cells were washed with ice cold PBS and incubated with acid-wash buffer (0.2 M glycine and 0.15 M NaCl, pH 3.0) for 5 min. Subsequently, cells were washed two times with PBS, and finally lysed in RIPA supplemented with cOmplete protease inhibitor cocktail and PhosSTOP phosphatase inhibitors.

### Tissue preparation and immunofluorescence

Whole mouse organs and crural muscles were frozen immediately post-euthanasia. For protein isolation, the samples were snap frozen in liquid nitrogen. For cryosectioning, specimens were coated with Tissue-Tek O.C.T. compound (Sakura, The Netherlands) and frozen in ice-cold isopentane (∼-150°C). Human tissue samples from the marginal border of resected specimens were treated similarly during the first five hours after resection.

For immunofluorescence analysis tissues were cut into 10 μm sections (SLEE Cryostat, Germany), thaw-mounted on glass slides (Superfrost, Thermo Fisher Scientific Massachusetts, USA) and air-dried for 1 h at 37 C. Cryosections were immersed for 4 min in ice-cold acetone and again air-dried for 1 h at RT. Specimens were transferred in humidified chamber, rehydrated with phosphate-buffered saline (PBS) for 10 min continued with permeabilization using 0.1% Triton X-100 plus 0.05% Tween-20 in PBS for either 15 min. Samples were then incubated for 60 min at RT in blocking buffer (5% normal donkey serum (Dako) plus 0.05% Tween-20 in PBS) and incubated overnight at 4°C in a humidified chamber with primary antibodies against human- or mouse-asprosin (polyclonal rabbit, dilution 1:60, lab-made). After washing 3 times for 5 min in PBS containing 0.05% Tween-20, specimens were incubated with secondary antibodies (Alexa Fluor 488-conjugated donkey anti-Rabbit IgG (H+L) # R37118, ThermoScientific, Massachusetts, USA, dilution 1:300). In the presence of DAPI (1μg/mL) (#62248, Thermo Fisher Scientific, Massachusetts, USA) for 1h at RT. Primary and secondary antibodies were diluted into Antibody Dilution Buffer (#AL120R100, DCS Diagnostics, Germany). Finally, specimens were washed once with PBS containing 0.05% Tween-20 for 5 min then twice with PBS (5 min each) and mounted with ProLong Gold Antifade Mountant (#P10144, Thermo Fisher Scientific Massachusetts, USA). Sections processed with the only secondary antibodies served as controls to exclude autofluorescence and non-specific binding. cOmplete protease inhibitor cocktail (one tablet per 50 ml PBS, cOmplete™ ULTRA Tablets, Mini, EDTA-free, Roche) was added to all solutions and buffers used for previously described procedures, except for the permeabilization solution.

### Cell culture

All cell lines used in the study were provided from American Type Culture Collection (ATCC). HDF (primary human dermal fibroblasts) from control as well as HCH (primary human chondrocytes) were established from biopsies. Cells were cultured in DMEM with 10% fetal calf serum (FCS) and 1% penicillin/ streptomycin, they were grown in incubators maintained at 37°C and 5% CO2. Cells were fed twice per week and split regularly.

### Surface plasmon resonance

SPR experiments were performed as described previously (10, 13) using a BIAcore 2000 system (BIAcore AB, Uppsala, Sweden). Recombinant human asprosin (#761902, Biolegend, San Diego, CA, USA) was immobilized at 1000 RUs to a CM5 sensor chip using the amine coupling kit following the manufacturer’s instructions (Cytiva, Uppsala, Sweden). Interaction studies were performed by injecting 0-320 nM recombinant fibrillin-1 fragments (Start-EGF, or rF90), or fibronectin in HBS-EP buffer (0.01 M HEPES, pH 7.4, 0.15 M NaCl, 3 mM EDTA, 0.005% (v/v) surfactant P20) (Cytiva, Uppsala, Sweden). Kinetic constants were calculated by nonlinear fitting (1:1 interaction model with mass transfer) to the association and dissociation curves according to the manufacturer’s instructions (BIAevaluation version 3.0 software). Apparent equilibrium dissociation constants (KD values) were then calculated as the ratio of kd/ka.

### Crosslinking by transglutaminase 2 (TG2) and fluorescent-monodansylcadaverine

20 μg of purified asprosin (5 μg/μl) was buffer exchanged by Amicon Ultra Centrifugal Filters (cut off: 3 kDa), with PBS containing 5 mM CaCl2 and treated with 2 μg transglutaminase 2 (#T5398, from guinea pig liver, Sigma-Aldrich, Darmstadt, Germany) (enzyme: substrate ratio, 1:10 (w/w)) in a total volume of 300 μl. The reaction mixture was incubated at 37°C. 15 μl of the reaction mixture was withdrawn after 5, 10, and 15 min. The reaction was directly stopped by adding 5 μl of Laemmli Buffer and the samples were analyzed by SDS-PAGE and western blot. For the control experiment tTG was omitted or samples were treated with 0.25 M EDTA to sequester calcium ions and inhibit tTG activity. Fluorescent-monodansylcadaverine (MDC) (# D4008, Aldrich, Darmstadt, Germany) was used as amine donor and acceptor sites for tissue transglutaminase enzyme activity detection as previously suggested and described (52). MDC was dissolved in DMSO, 2 mg in 1.5 ml DMSO to obtain a stock solution of 4 mM MDC. 40 μg of purified asprosin (5 μg/μl) was buffered exchanged with PBS containing 5 mM CaCl2 and 40 μM MDC in a total volume of 120 μl. the reaction mixture was then treated with 4 μg tTG (1 μg/μl) and incubated at 37°C. 15 μl of the reaction mixture was withdrawn after 5, 10, 15, 20, 30, and 45 min, then the reaction was directly stopped by adding 5 μl Laemmli Buffer subjected to SDS-PAGE. The gel was then fixed with 25% isopropanol, 10% acetic acid and photographed on a 300 nm UV, then subjected to Coomassie Blue Staining.

### DSS (disuccinimidyl suberate) crosslinking

DSS (#21655, Thermo Fisher Scientific Massachusetts, USA) is a water-insoluble chemical crosslinker and was dissolved in dimethyl sulfoxide (DMSO), 1 mg in 55.29 μl DMSO to obtain 800 μM DSS, then serially diluted (1:2) into DMSO to obtain following DSS concentrations 400, 200, and 100 μM DSS. Cross-linking reactions were conducted in a total volume of 15 μl containing 1 μg asprosin (0.5μg/μl), 1.5 μl of freshly prepared DSS to obtain final DSS concentrations (80, 40, 20, and 10 μM) and complete to the 15 μl final volume with PBS. The reaction mixtures were incubated at RT for 30 min was directly stopped by adding 5 μl of Laemmli Buffer and the samples were analyzed by SDS-PAGE and western blot. For the control experiment, 1.5 μl of DMSO was used instead of DSS.

### Preparation of fluorescently labelled asprosin (Asprosin-550)

1 mg of ATTO 550 dye (#AD 550-31, ATTO-TEC GmbH, Germany) was freshly prepared by dissolving it in 100 μl DMSO directly before coupling to asprosin. 500 μg of purified human asprosin was buffered exchanged into a coupling buffer (PBS containing 10 mM sodium bicarbonate and 100 mM sodium hydroxide, pH 8.3) by Amicon Ultra Centrifugal Filters (cut-off: 3 kDa). The coupling reaction mixture was prepared by mixing the dissolved ATTO 550 dye and the buffer-exchanged asprosin in the coupling buffer to a final volume of 5 ml. The mixture was then incubated for 1 h, protected from light at RT. The coupling reaction mixture was concentrated by using Amicon Ultra Centrifugal Filters (cut-off: 10 kDa), (starting volume: 5 ml, end volume: 0.3 ml). The concentration of fluorescently labelled asprosin (Asprosin-550) was measured by NanoDrop One (Thermo Fisher Scientific, Massachusetts, USA). The quantification was performed by measuring the protein absorbance at 280 nm and the ATTO-550 dye absorbance at 550 nm. The integrity of the conjugated asprosin was additionally analyzed by SDS-PAGE and subjected to Coomassie Blue Staining (supplementary Fig. S1B). The fluorescence of the conjugated asprosin was visualized by ChemoStar Touch ECL & Fluorescence Imager (Intas Science Imaging Instruments GmbH, Göttingen, Germany).

### UNcle analysis

Samples were diluted into 5× buffers resulting in a final concentration of 100 mM buffer and 150 mM NaCl. 2 μl of protein, 2 μl of buffer and 6 μl of water were used for each sample. 16 buffer conditions were loaded via glass capillaries into the UNcle instrument. Dynamic light scattering measurements were performed at a constant temperature of 25°C with attenuation set to automatic. Reported values represent the average of three readings per condition. The melting and aggregation onset measurements were performed by heating the sample to 95°C at 1°C intervals and 60 sec incubation at each temperature. The instrument acquires a full spectrum of tryptophan fluorescence using a 266 nm laser to excite and scan between 250 and 550 nm and monitors the level of scattering of both the 266 nm and 473 nm lasers. Tagg is the aggregation onset temperatures defined as the beginning of the aggregation curve, whilst the melting temperature (Tm) is defined as the mid-point in the melting curve, which represents the level of tryptophan exposure to solvent.

### Asprosin structure prediction

The structure of asprosin (fibrillin-1 P3555 residues 2732-2871) was predicted in Alphafold2 using colabfold (DOI: 10.1038/s41592-022-01488-1).

## Acknowledgments

We would like to thank the CECAD Cologne Proteomics Facility for conducting mass spectrometry analysis. Funding for this study was provided by the Deutsche Forschungsgemeinschaft (DFG, German Research Foundation) project ID 384170921: FOR2722/ B1 to M.P., and FOR2722/ M2 to E.H.R., and G.S. B.C. is a senior clinical investigator of the Research Foundation Flanders. This work was supported by a grants of the Special Research Fund of Ghent University (grant 01N04516C and BOF21/GOA/019 to B.C.). The Ghent University Hospital is a member of the European Reference Network for Skin Disorders (ERN-Skin).

## Author contributions

Y. A. T. Morcos, C. Baldock, and G. Sengle and designed research. Y. A. T. Morcos, G. Pryymachuk, A. R. F. Godwin, S. Lütke, T. A. Jowitt, T. Hoffmann, A. Tenbieg, and N. Piekarek performed research. Y. A. T. Morcos, G. Pryymachuk, A. R. F. Godwin, T. A. Jowitt, S. Lütke, and C. Baldock analyzed data. W. Bloch, B. Callewaert, U. Drebber, O. Grisk, A. Niehoff, M. Odenthal, and Y. Ladilov provided essential reagents. M. Paulsson provided expert advice. Y. A. T. Morcos, and G. Sengle wrote the manuscript.

