## supplemental Figures for "Transglutaminase mediated asprosin oligomerization allows its tissue storage as fibers"

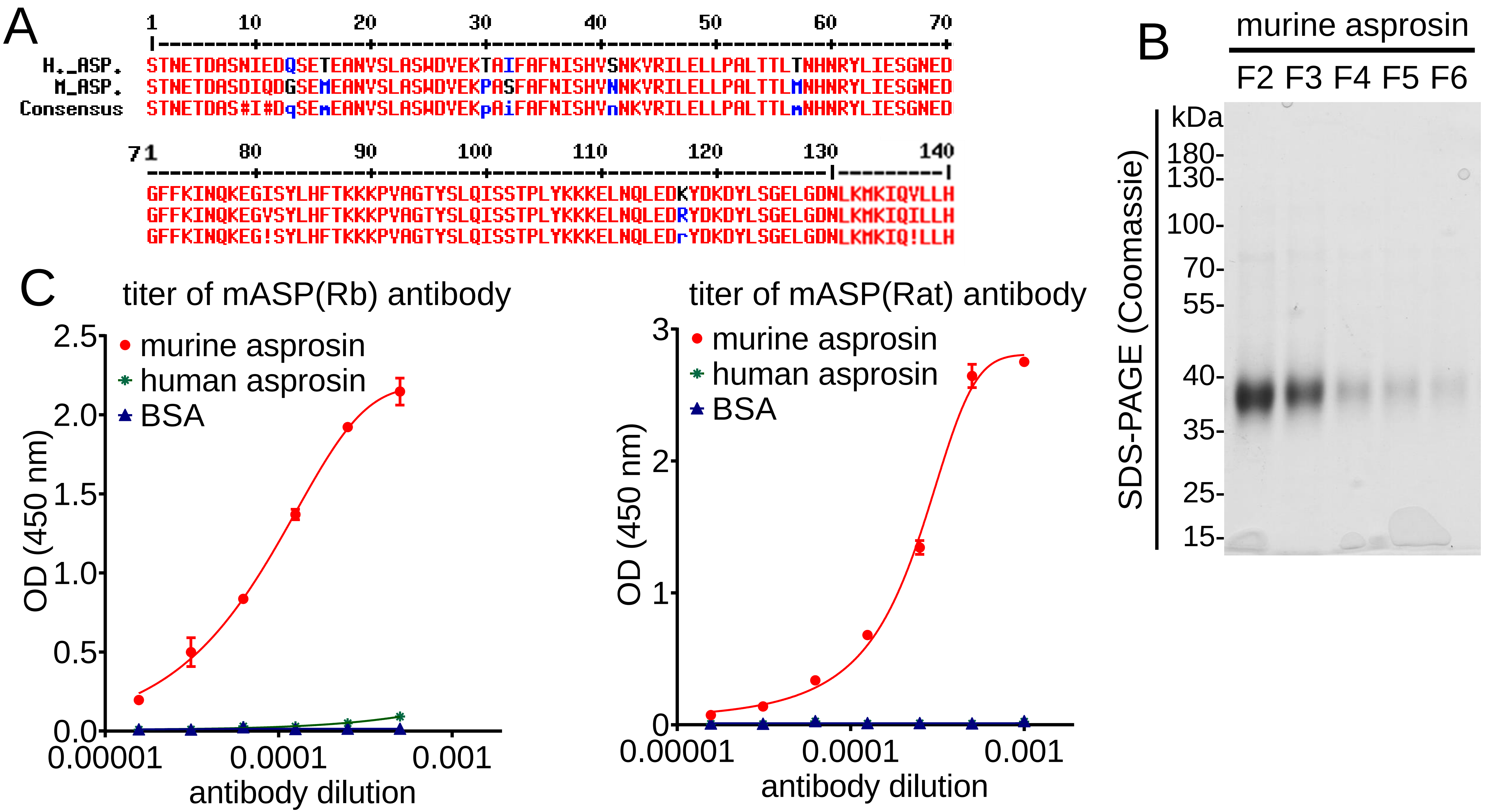


**Supplementary Figure S1: Generation and evaluation of polyclonal anti murine asprosin antibodies. A.** Sequence alignment between human and murine asprosin using Multalin software showed 92.14 % sequence identity between both proteins. Red color represents highly conserved residues; blue color represents weakly conserved residues; symbols:! represents either I or V; # represents any one residue of NDQEBZ; i represents either I or L or T. **B.** Coomassie stained quality control gel of recombinantly produced and purified murine asprosin showing >95 % purity after affinity chromatography. **C.** Affinity purified polyclonal antibodies raised in rabbit (left) and rat (right) showed high specificity in detecting coated murine asprosin (100 ng/well) by direct ELISA with no crossreactivity to human asprosin or BSA. Data points represent mean ± SD of duplicates.

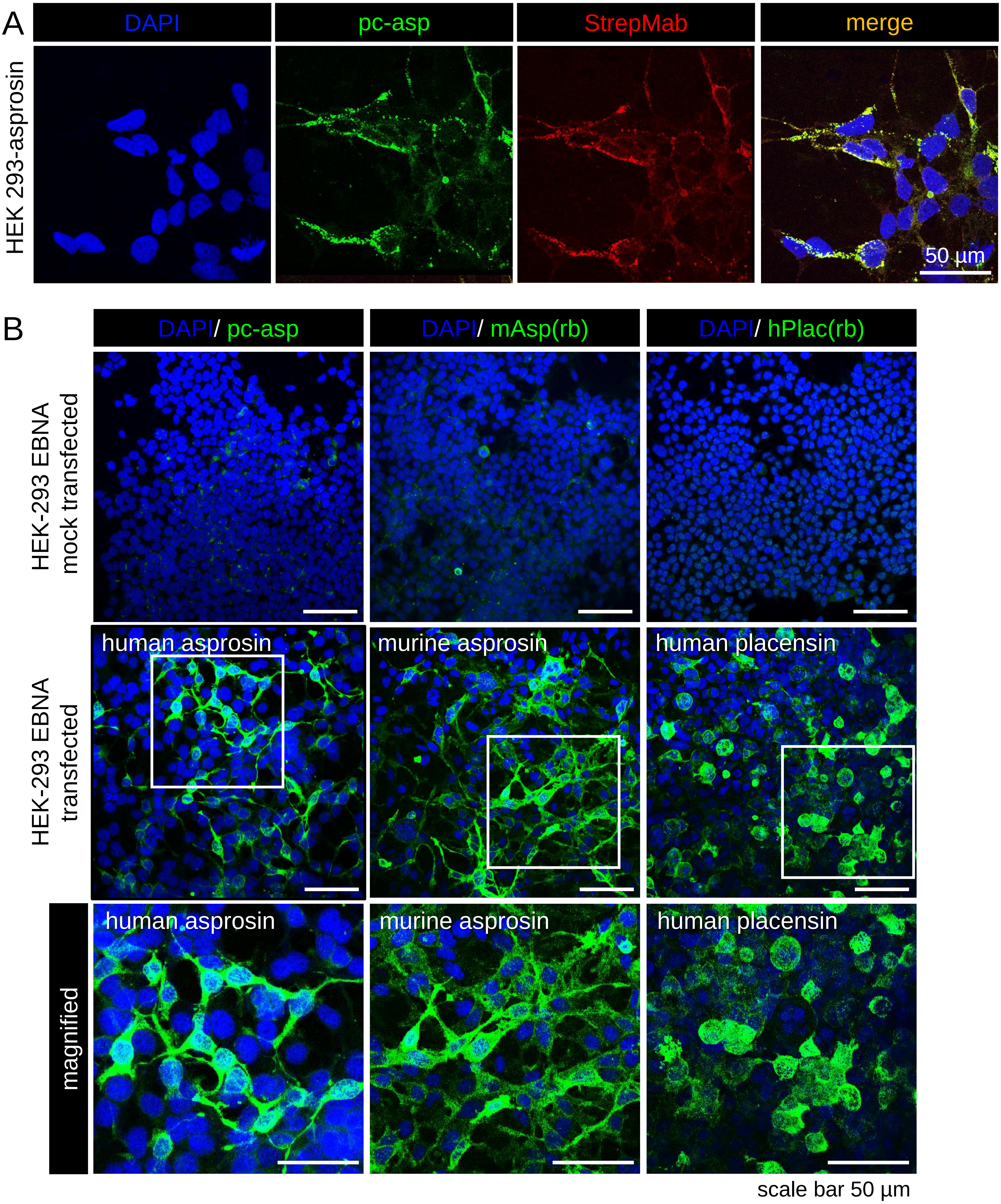


**Supplementary Figure S2: Immunofluorescence evaluation of generated labmade asprosin and placensin antibodies. A.** Immunofluorescence analysis of HEK-293 cells transfected with human asprosin-2×Strep-tag II using pc-asp (green), StrepMAB Ab (red), and DAPI (blue, nuclei) demonstrates co-localization of signals from pc-asp and StrepMab antibodies. **B.** Comparative immunostaining examination of the generated labmade human asprosin (pc-asp), mouse asprosin (mASP(rb) Ab), and human placensin (hPlac(rb) Ab) antibodies (green) in HEK-293 EBNA mock transfected cells and cells transfected with a construct expressing the indicated protein. (top panel) No signal was detected in mock transfected cells. (middle and bottom panel) Positive staining was detected by antibodies in human asprosin, mouse asprosin, or human placensin transfected cells confirming their specificity in immunofluorescence application. Two-fold magnified areas are marked by white boxes. Images were obtained from a Leica SP8 confocal microscope and were processed using Leica Application Suite X (LAS X) software version 3.7.5.2. Fiji/ImageJ (version 1.53t) software was used to obtain average intensity Z-projection.

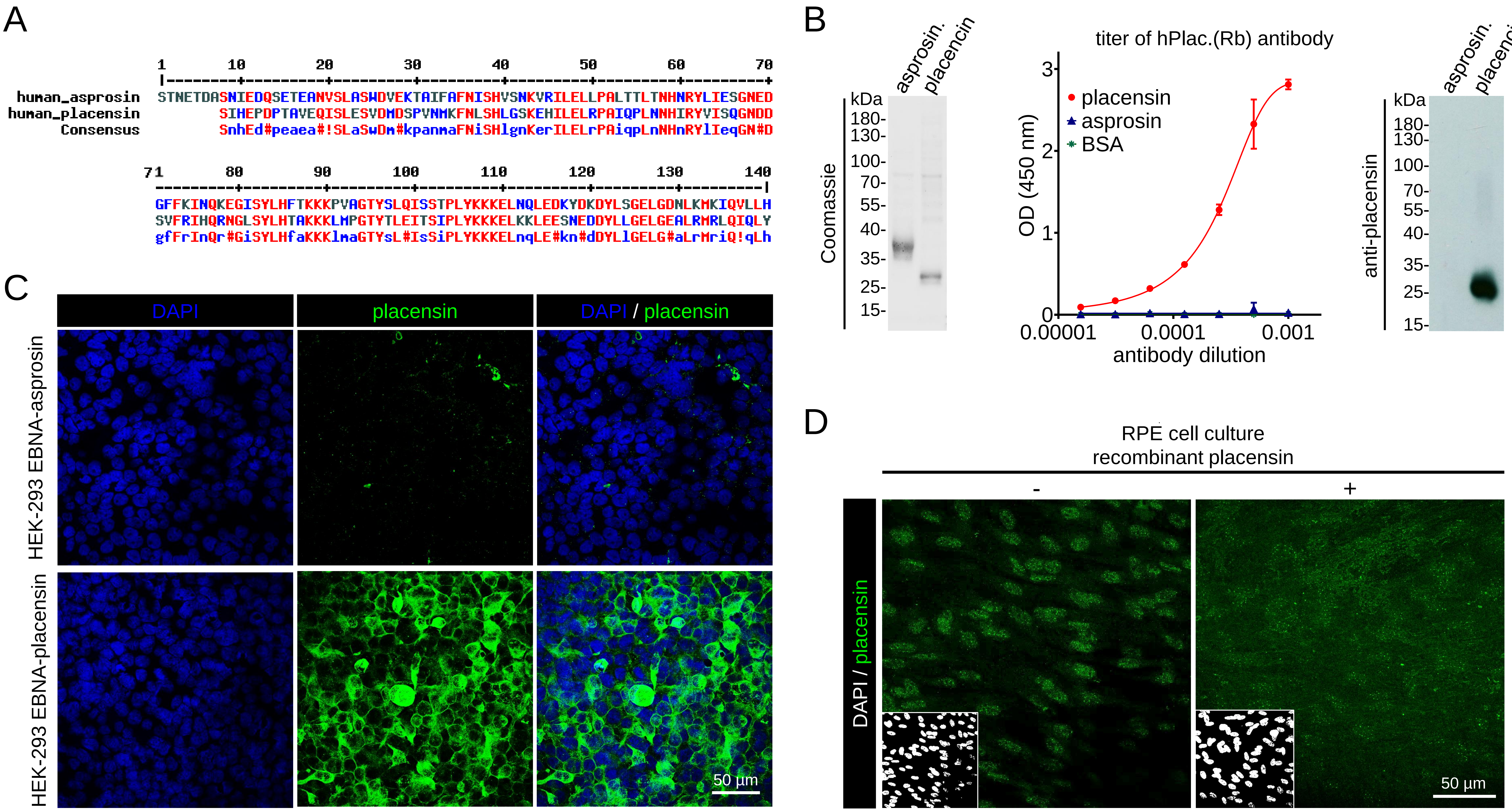


**Supplementary Figure S3: Administration of placensin to RPE cells failed to form fibers. A.** Primary sequence alignment between human asprosin and human placensin performed by Multalin software program reveals 47% identity and 67% similarity between the two proteins. Red color represents highly conserved residue; Blue color represents weakly conserved residue; Symbols:! represents either I or V;# represents any one residue of NDQEBZ; i represents either I or L or T. **B.** (left) Coomassie stained quality control gel of recombinant produced and purified (>95% purity) asprosin and placensin after affinity chromatography. (middle) Affinity purified polyclonal human-placensin antibody raised in rabbit, hPlac.(Rb) antibody showed high specificity in detecting coated placensin (100 ng / well) by ELISA. No cross reactivity to aprosin or BSA was detected indicating the specificity of the antibody. Data points represent mean ± SD of duplicates. (right) Human placensin antibody specifically recognizes human placensin in western blot analysis, with no observable cross reactivity to asprosin. **C.** Immunofluorescence analysis of HEK-293 cells transfected with (top) human asprosin (N-His$_6$) and (bottom) human placensin (C-2×Strep-tag II) with hPlac.(Rb) antibody (green) and DAPI (blue, nuclei) shows specific intracellular placensin staining, with no cross reactivity to asprosin. **D.** Recombinant placensin (5µg/ml) was administered to RPE cells. 48 h after placensin treatment, cells were incubated with hPlac.(Rb) antibody (green) and DAPI (blue, nuclei). Placensin immunostaining showed only intracellular placensin signals and no presence of fibers. Images were obtained from a Leica SP8 confocal microscope and were processed using Leica Application Suite X (LAS X) software version 3.7.5.2. Fiji/ImageJ (version 1.53t) software was used to obtain average intensity Z-projection.

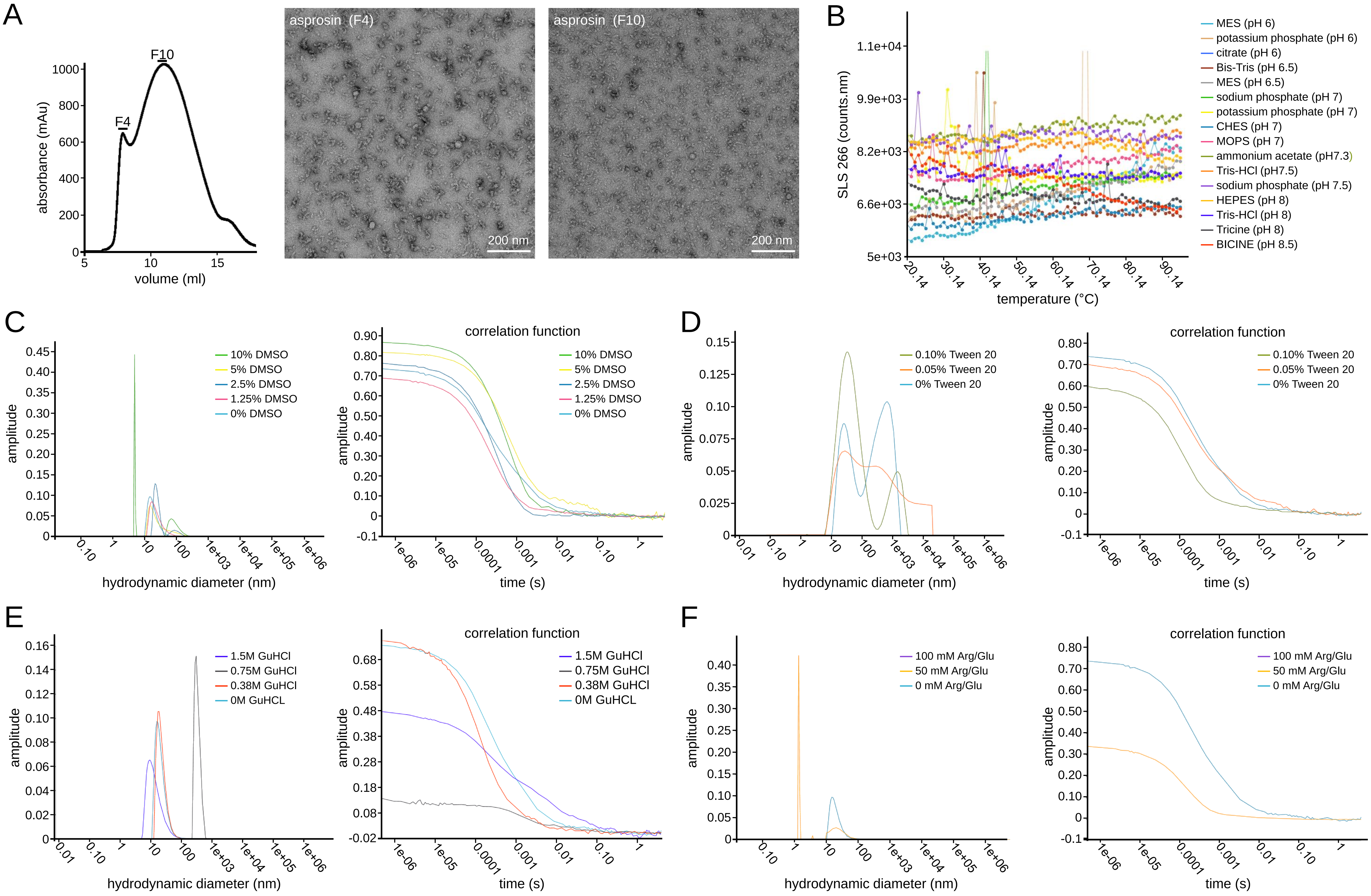


**Supplementary Figure S4: Analysis of asprosin multimers using SEC, TEM and UNcle analyzer. A.** Recombinant asprosin subjected to SEC analysis and cryo-EM visualization of fractions 4 and 10. The overview EM images show asprosin multimers in different shapes and sizes. **B.** Examination of asprosin thermostability in several buffers at temperature ranges from 20°C to 38°C shows no significant changes in the measured static light scattering (SLS) indicating no variations in the asprosin multimeric state. **C – F.** Measurement of hydrodynamic diameters and correlation functions of asprosin in presence of different buffer showing several degrees of asprosin dissociation in the indicated buffers.

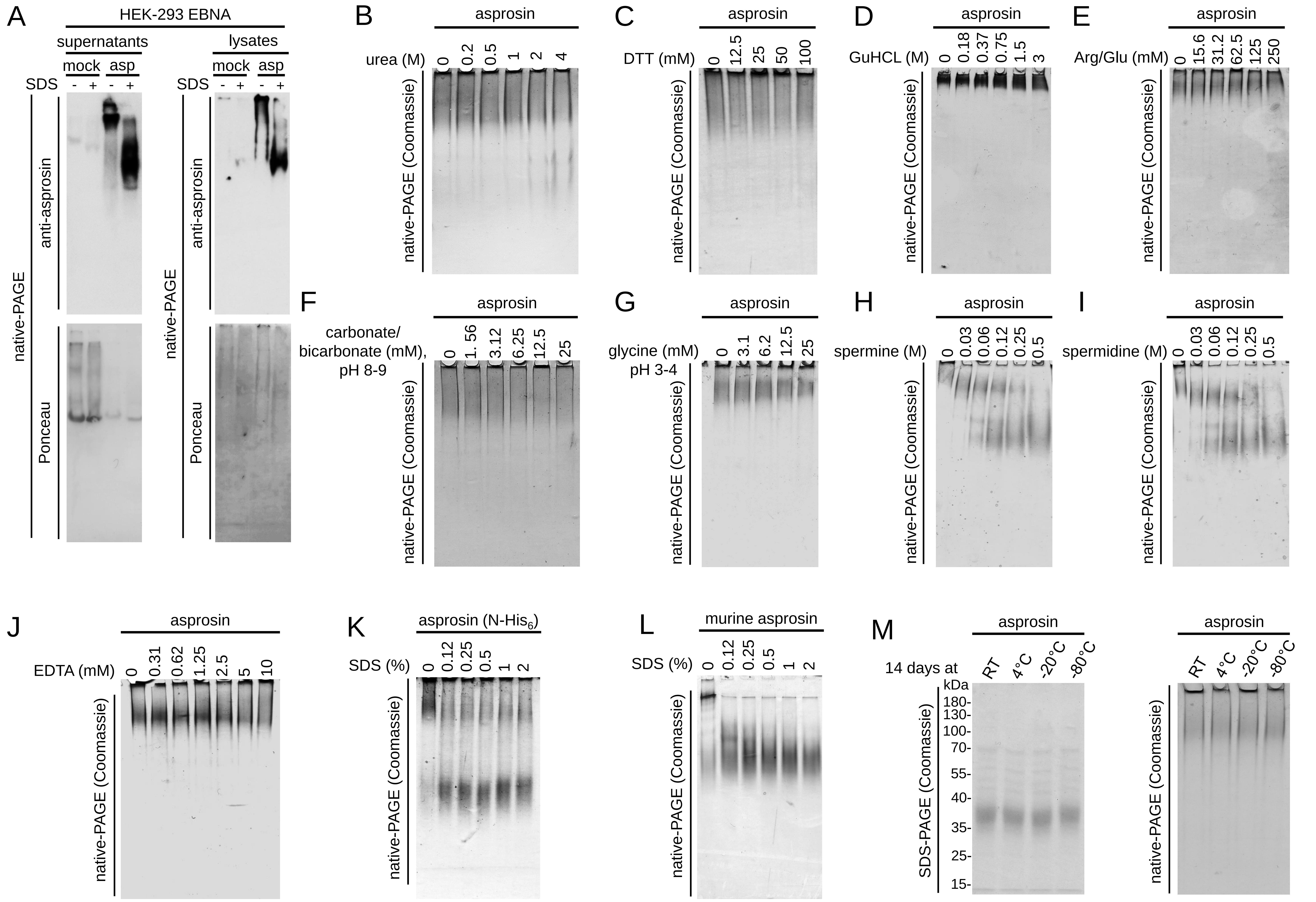


**Supplementary Figure S5: Impact of pH change and denaturing agents on asproisn multimerization A.** Western blot analysis of cell culture supernatants (left) and cell lysates (right) from stably transfected HEK-293 EBNA asprosin overexpressing cells. Supernatants and cell lysates of mock and asprosin transfected cells were treated with 1% SDS and subjected to 10% native-PAGE followed by western blot analysis. Immunoblots show the dissociation of asprosin oligomers upon SDS treatment. **B.** Diluting asprosin in urea solutions ranging from 0.2M to 4M urea followed by native-PAGE analysis shows mild dissociation of asprosin oligomers in 4M Urea. **C.** Addition of dithiothreitol (DTT) up to 100 mM shows no effect on asprosin multimerization. **D.** Diluting asprosin in GuHCL solutions ranging from 0.18M to 3M then subjected to native-PAGE analysis shows no dissociation of asprosin oligomers. **E.** Treating asprosin with a serial concentration of arginine/glutamine ranging from 15.6mM to 250mM shows no dissociation of asprosin oligomers. **F.** Effect of basic pH on the multimeric state of asprosin in carbonate/bicarbonate solution (pH 8 – 9) reveals no detectable change compared to the untreated sample. **G.** Concentration-dependent increase of asprosin oligomerization in acidic conditions upon buffer exchange in glycine solution (3 – 25 mM, pH 3 – 4). Native-PAGE analysis indicates aggregation of asprosin n of asprosin multimers. **H-I.** Addition of spermine and spermidine on asprosin multimerization shows dissociation of asprosin oligomers similar to SDS treatment (Fig. 4B, middle). **J.** Treatment of asprosin with EDTA shows no detectable effect on its multimeric state. **K.** Native-PAGE analysis of untreated and SDS-treated asprosin (N-$His_6$) shows dissociation of asprosin. **L.** Analysis of murine asprosin after and before SDS treatment shows similar pattern as human asprosin (Fig. 4B, middle). **M.** Coomassie staining of native and SDS-PAGE analyses of affinity purified asprosin after incubation at RT, 4°C, -20°C and -80°C for 14 days revealed no changes over time and temperature. For each experiment 3 µg of purified asprosin were used, samples in B and C were incubated for 10 min at 75°C before loading, and samples in D, E, F and G were incubated for 10 min at 37°C before loading.

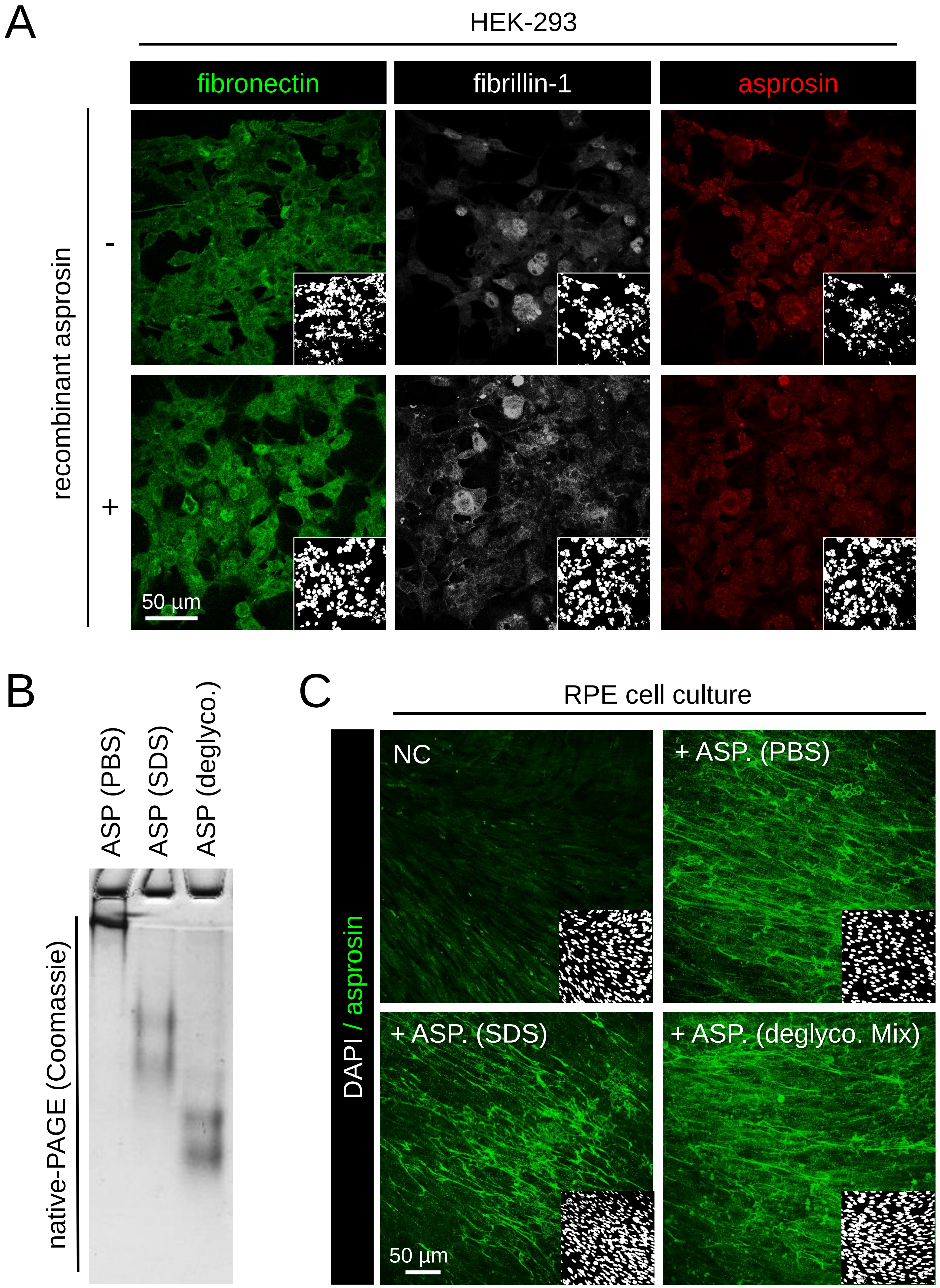


**Supplementary Figure S6: A. Impact of asprosin multimerization and glycosylation on fiber formation. A.** Immunofluorescence analysis revealed the absence of an intact fibrillin-1 and fibronectin fiber network in cultures of HEK-293 EBNA cells. Asprosin immunostaining showed no significant differences in staining patterns before and after asprosin treatment. **B.** Coomassie stained native-PAGE gel of recombinant asprosin after treatment with SDS or incubation in a deglycosylation mix showing a change in its migration after treatment. **C.** Asprosin positive fibers were observed after administration of different batches of asprosin (analyzed in B) to RPE cells. Images were obtained from a Leica SP8 confocal microscope and were processed using Leica Application Suite X (LAS X) software (version 3.7.5.2) and Fiji/ImageJ software (version 1.53t) to obtain average intensity Z-projection. Scale bars: 50 µm.

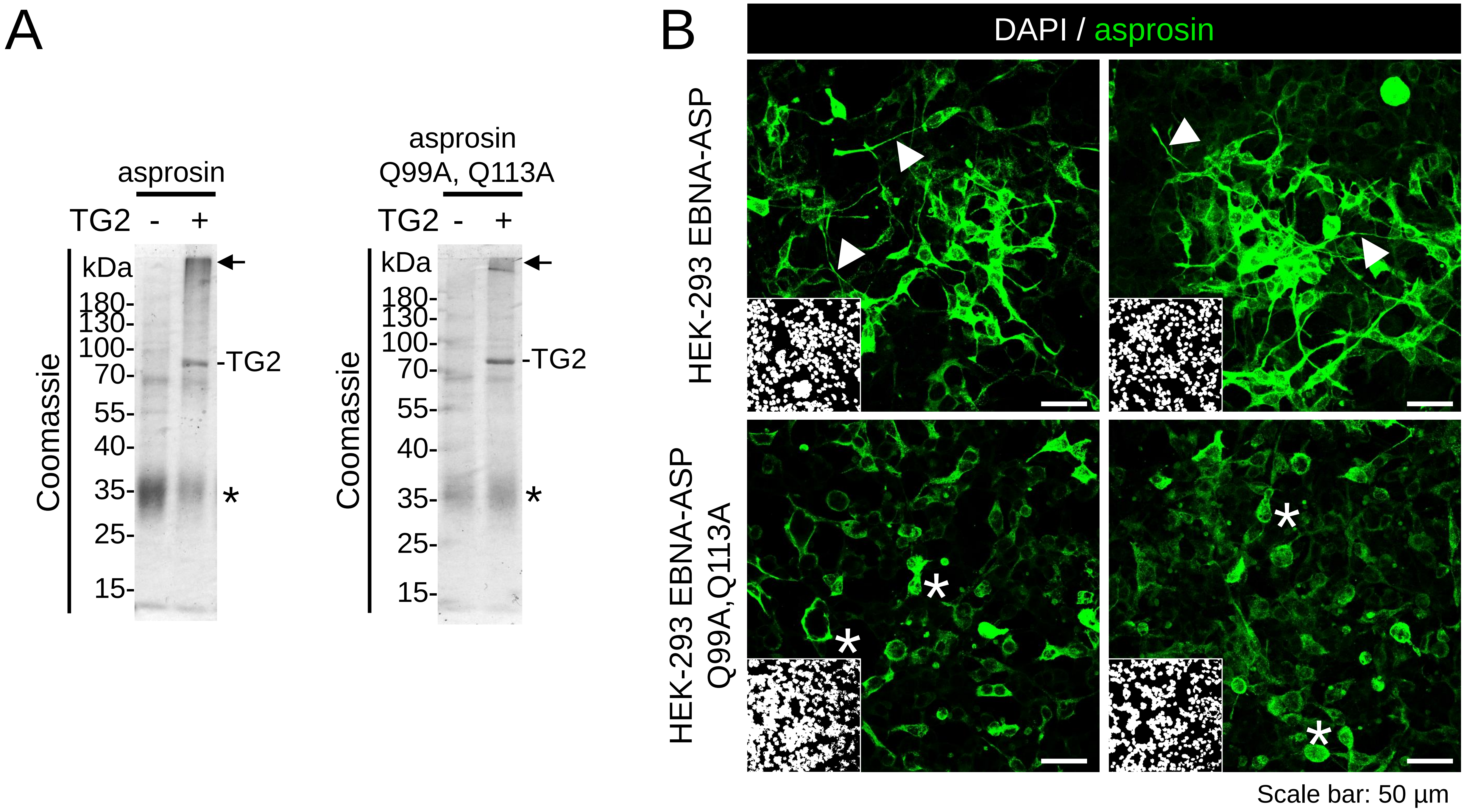


**Supplementary Figure S7: Mutant asprosin does not serve as a substrate for transglutaminase-mediated fiber formation. A.** Incubation of recombinant wild-type asprosin (left) and recombinant mutant asprosin (Q99A, Q113A, sequence shown in Fig. 10A) (right) in presence of TG2 results in a significant increase of a high-molecular weight band in wild-type asprosin compared to mutant asprosin (black arrows). Also TG2 addition resulted only in a depletion of monomeric wild-type asprosin (black asterisks). **B.** Immunofluorescence analysis using pc-asp Ab (green) and DAPI (white, nuclei) in HEK-293 EBNA cells overexpressing wild-type and mutant asprosin proteins. (top panel) Representative images from two independent experiments of HEK-293 EBNA cells transfected with a wild-type asprosin construct resulting in the formation of asprosin positive fibers (white arrowheads). (bottom panel) Representative images from two independent experiments of HEK-293 EBNA cells transfected with a mutant asprosin construct show only intracellular diffused asprosin signals. Images were obtained from a Leica SP8 confocal microscope and were processed using Leica Application Suite X (LAS X) software (version 3.7.5.2) and Fiji/ImageJ software (version 1.53t) to obtain average intensity Z-projection. Scale bars: 50 µm.

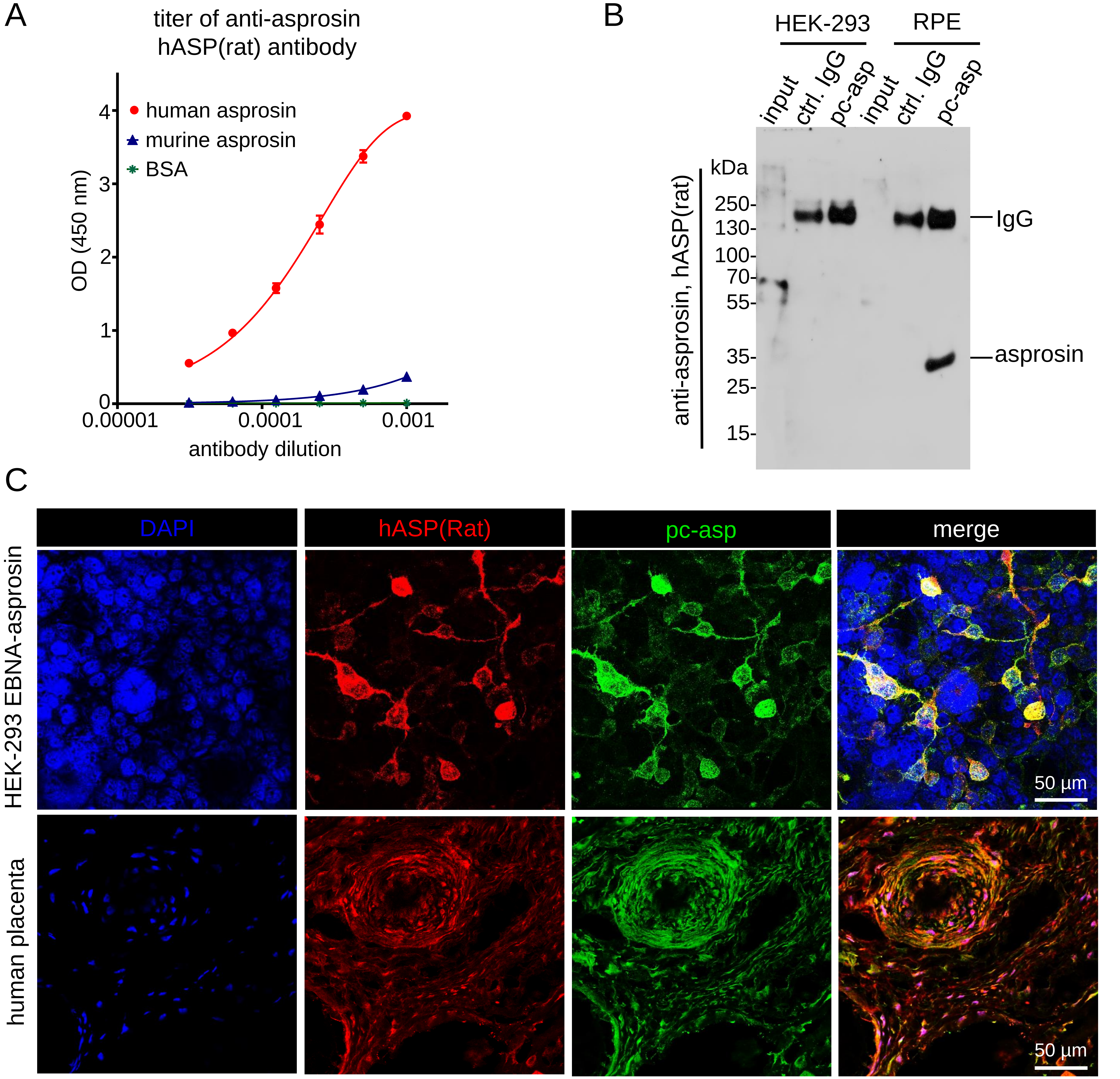


**Supplementary Figure S8: Generation of anti-asprosin hASP(Rat) antibody and its application in immunoprecipitation and immunostaining. A.** Affinity purified polyclonal anti-human-asprosin antibody raised in rat hASP(Rat) shows high specificity in detecting coated human asprosin (100 ng / well) by ELISA. No cross-reactivity to murine asprosin or BSA was detected. Data points represent mean ± SD of duplicates. **B.** Western blot analysis with hASP(Rat) ab of cell culture supernatants (input) and elution fraction after immunoprecipitation with rabbit control IgG (#26102, Thermo Fisher Scientific) (ctrl. IgG) and pc-asp antibody. **C.** Immunodetection of asprosin in cultures of asprosin overexpressing HEK-293 EBNA cells and in human placenta tissue using pc-asp (green), hASP(Rat) ab (red), and dapi (blue, nuclei) showing colocalization of the signals developed from both antibodies. Images were obtained from a Leica SP8 confocal microscope and. Images were processed using Leica LAS AF Lite 4.0 software and Fiji/ImageJ software to obtain average intensity Z-projection.

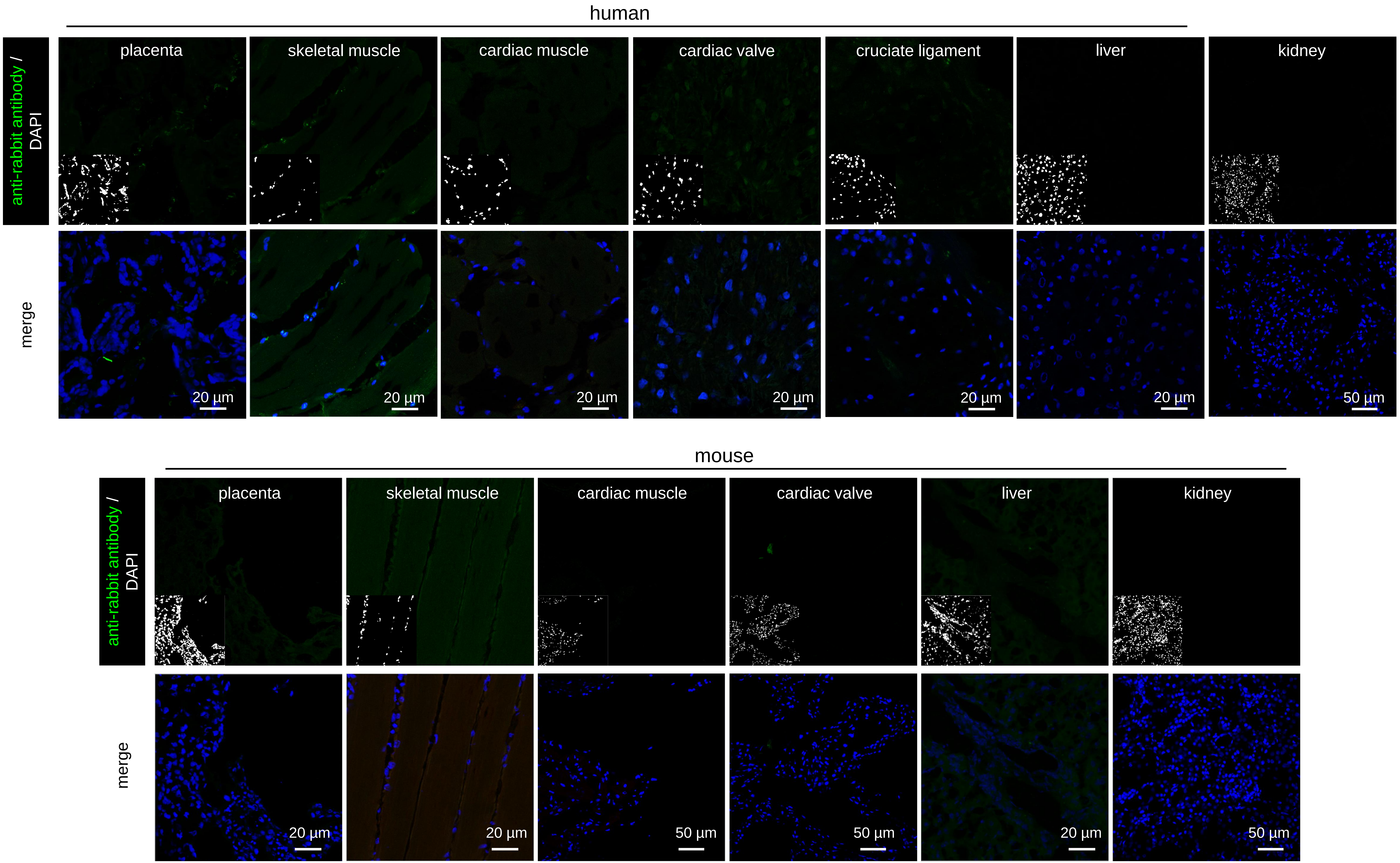


**Supplementary Figure S9: Negative immunofluorescence controls of human and mouse specimens**. Images show sections incubated with secondary antibody only (negative control) and DAPI. Images were obtained from a Leica SP8 confocal microscope and were processed using Leica LAS AF Lite 4.0 software and Fiji/ImageJ software to obtain average intensity Z-projection.
